# Chitin synthase targeting antifungal agents for Mucormycosis (Black fungus disease) caused by *Rhizopus delemar* - An *in-silico* study

**DOI:** 10.1101/2024.03.26.586901

**Authors:** Abhilash J George, Achuthsankar S Nair, B Vijayalakshmio

## Abstract

Mucormycosis, a severe fungal infection caused by *Mucorales* fungi, particularly *Rhizopus delemar*, has prompted the development of new and more potent antifungal drugs due to the emergence of drug-resistant strains. Currently used drugs are known to be toxic to human cells also, which is a major drawback in their administration. Antifungal drugs commercially under use and their structurally modified analogues along with 30 plants reported with antifungal activity from IMPPAT database, totalling 229 compounds, were screened against the best modelled structure of Chitin Synthase, a key target enzyme of *Rhizopus delemar* by *in-silico* molecular docking studies. Computerized methods to assess absorption, distribution, metabolism, excretion, and toxicity (ADMET) profiles was employed to further filter key lead drug candidates from among the docked ligands that could act as potential antifungal medications. Modified structures of Posaconazole, Nikkomycin and Isavuconazole were found to give better docking results than compared to the original drugs, though they failed to comply with ADMET parameters. Among the phytochemicals, 1-nonacosanol displayed binding affinity (highest LibDock score) greater than the synthetic drugs and their derivatives, though it too failed to comply with ADMET parameters. The phytochemical Dihydrocapsaicin was found to be the best compound that satisfied all the parameters such as ADMET, TOPKAT, Lipinski’s and Veber’s rule along with good binding affinity. All in all, this novel study paves a way to investigate the interaction of Chitin Synthase inhibitors at molecular level to achieve an alternative solution with less toxic effects to treat Mucormycosis.

## INTRODUCTION

Mucormycosis is an uncommon disease often associated with various underlying infections that weaken the immune system of those infected. The primary risk factors for this invasive fungal infection include intense chemotherapy, immune-compromising conditions like HIV, and the use of immunosuppressive medications (Shahi and Mishra, 2021). It is a serious fungal infection caused by *Mucorales*, with *Rhizopus delemar* being the most aggressive causative agent. Other fungal organisms such as *Lichtheimia, Cunninghamella, Mortierella, and Saksenaea* can also be responsible, albeit less frequently. The disease spreads through mold spores, commonly through inhalation, contaminated food, or direct contact with open wounds (trauma).

Symptoms vary depending on the affected body part and may include a runny nose, darkened skin areas, facial swelling, headaches, fever, cough, blurred vision, and more. Diagnosis involves biopsy, culture, and medical imaging to determine the extent of the infection. Treatment for Mucormycosis typically involves the use of Amphotericin-B and Isavuconazole, either alone or in combination with other antifungal medications (Sipsas et al., 2018). Invasive cases may require surgical intervention, which can be quite extensive. In situations where the infection affects the nasal cavity and the brain, the removal of infected brain tissue may be necessary. Surgical procedures involving the removal of the palate, nasal cavity, or eye structures can lead to significant disfigurement, and multiple operations may be required in some instances.

Moreover, the alarming nature of Mucormycosis lies in its remarkably fast rate of spread. A mere 12-hour delay in diagnosing the condition can prove fatal, and historically, half of all cases of Mucormycosis have been detected postmortem, intensifying the gravity of the situation (Singh et al., 2023).

A disproportionately high number of black fungus cases were reported in India as post-COVID infections shortly after the COVID-19 ‘[’ wave. In India, the incidence was 80 times higher than in wealthy countries. It was discovered that incidences were comparatively high among patients who took corticosteroids, including those admitted to intensive care units and those who were in the hospital for extended periods of time. Also, having diabetes increases one’s likelihood of developing a black fungus infection (and more Indians have diabetes than people in Western countries). The associated disorders affect the Mucormycosis mortality rate, which is greater than 50%. Patients with disseminated illness, infections of the central nervous system, or protracted neutropenia have a mortality rate of 90–100% (Jha et al., 2022). Sadly, the need for innovative therapeutic medications to treat the condition is urgent because of the unacceptable high mortality rate, restricted treatment alternatives (which have associated toxicities), high expense of managing Mucormycosis, and the very disfiguring surgical procedures, among other factors.

As discussed above, one of the main problems associated with treating of Mucormycosis is the non-availability of effective treatment methods. Most of them involve surgery and removal of infected part(s), which is painful as well as economically expensive (Chaudhary, Tupe and Deshpande, 2013). Many of the potential targets of this fungus are undiscovered, as large number of studies have not been conducted on this fungus yet. It is surprising to notice that, out of the limited number of studies, more than 90% of studies are from India, with most publications in the year 2022. Most of the antifungals available for this fungus targets cell membrane, and the associated sterols. This is a matter of concern as fungal cell membranes are similar to the human cellular membranes, and hence, this causes toxicity to human cells also.

### AIM AND OBJECTIVES

It was hypothesized that computer-aided screening of a library of phytochemicals (available in databases such as IMPPAT - Indian Medicinal Plants, Phytochemistry and Therapeutics) against the chitin synthase enzyme of *Rhizopus delemar* would help us to discover some of the lead molecules (phytochemicals with biological activity as well as fulfilling certain criteria) which can be further utilized to check for the real-world efficacy of these lead molecules (*in-vitro* and *in-vivo* studies) against the pathogen, to be administered in human subjects.

One of the main objectives of this project was to partially validate the phytochemicals present in the database under consideration which show appropriate properties suitable to be called as a drug candidate, with the help of computerized tools and software in order to simplify the process of drug discovery, in terms of time, money, labour, etc., which usually takes more than 15 years to be completed from scratch. The project would be completed with creating of a pipeline for partially validating phytochemicals against Chitin Synthase of *R. delemar*.

Apart from this, another major objective of this project was to check for the properties and binding efficacy of modified drugs (or drug analogues) derived from conventionally used antifungals against this disease such as Amphotericin B, Posaconazole, Isavuconazole, Nikkomycin, and Polyoxin. If these modified drug molecules were found to be more effective than their parent molecules, further validation can be performed to check for the real-world efficacy of these modified drugs.

### CHITIN SYNTHASE AS A SUITABLE TARGET

Chitin, a β-1,4 linked N-acetyl-D-glucosamine polymer, is an important constituent of the cell wall. Chitin content in the cell walls of fungus ranges from 2% in certain yeasts to 61% in some filamentous fungi. No matter the ratio, chitin seems to be necessary for fungal growth and survival.

In a transglycosylation reaction, which Chitin Synthase catalyses, sugar residues are transferred from UDP-GlcNAc to the lengthening chitin chain, releasing uridine diphosphate in the process. As chitin is absent in plants and animals, its synthesis is a promising target for development of antifungal agents (Chaudhary, Tupe and Deshpande, 2013).

The currently used drugs against Mucormycosis, as discussed earlier include Amphotericin B, Posaconazole, and Isavuconazole.

Amphotericin B is one of the most commonly used antifungal medication. Ergosterol, a substance present in the cell membrane of certain fungus, is where Amphotericin acts. It causes the development of ion channels once it undergoes complexation with ergosterol, which results in the loss of protons and monovalent cations. The depolarization impact finally results in the fungus cells being destroyed when combined with concentration-dependent processes. It also causes oxidative damage within cells, leading to the generation of free radicals and enhanced membrane permeability, in addition to its main method of action. Amphotericin B also has a stimulating impact on phagocytic cells, which helps to remove fungus infections (Noor and Preuss, 2023).

Apart from Amphotericin B, another important drug which is used for step-down therapy after administering Amphotericin B, is Posaconazole, which works by blocking the enzyme lanosterol 14α-demethylase, which leads to the inhibition of biosynthesis of ergosterol, which is a crucial part of the fungal cell membrane. As a consequence of this inhibition, a series of events take place, which include accumulation of methylated sterol precursors and depletion of ergosterol within the fungal cell membrane. Due to these events, the structure and function of the fungal cell membrane is destroyed and this leads to the antifungal activity of Posaconazole (Chen et al., 2020).

Isavuconazole is another drug which is used to treat patients infected with Black Fugus. This drug is also known to work on the same principles of Posaconazole (Ellsworth and Ostrosky-Zeichner, 2020).

Only Nikkomycin and Polyoxin, which were obtained from *Streptomyces* sp. culture filtrates, are recognised chitin synthase inhibitors at this time. Unfortunately, they have a very restricted spectrum and very little activity.

It is important to mention here that although Amphotericin B, Posaconazole and Isavuconazole target the sterol component of the fungal cell, these drugs can be used employed as control molecules in order to compare the results obtained after docking these modified drug molecules, derived from these drugs itself against the target protein of interest.

## METHODOLOGY

### Protein sequence retrieval

The amino-acid sequence for the Chitin Synthase (Accession: EIE85441) of *Rhizopus delemar* was derived from NCBI protein database. With the aid of protein structure prediction tools such as SWISS-MODEL, Phyre2 and Robetta, the retrieved sequence was used to create a 3D protein model, the steps for which are detailed in the following sections (Jha et al., 2022). It is worth mentioning here that all the different online tools used to model the protein structure work on different algorithms and parameters.

### Analysis of physicochemical properties of the protein

The ProtParam tool by ExPasy was utilized to calculate various physico-chemical properties of the protein, including Molecular weight, theoretical pI, Grand-average of hydropathy (GRAVY), half-life, aliphatic index (AI), instability index, and amino acid composition. The tool is accessible at ProtParam (https://web.expasy.org/protparam/) and enables the computation of physical and chemical parameters for a user-provided protein sequence or a protein sequence which is stored in SWISS-Prot or TrEMBL.

### Secondary structure prediction

To determine the secondary structure features of the protein, several methods can be utilized, including CFSSP (Chou and Fasman Secondary Structure Prediction) (Kumar, 2013). CFSSP, which employs the Chou-Fasman algorithm, analyzes the relative frequencies of each amino acid in α-helices, β-sheets, and turns based on known protein structures resolved with X-ray crystallography. The Chou and Fasman Secondary Structure Prediction Server is available at (https://www.biogem.org/tool/chou-fasman/).

Another approach for secondary structure prediction is PSI-Pred (PSI-blast based secondary structure prediction) (Banik et al., 2022). (Self-Optimized Prediction Method from Alignment) SOPMA is also a method that can be used for the determination of protein secondary structure (Madanagopal et al., 2022). These methods provide valuable insights into the secondary structure elements of proteins, including α-helices, β-turns, and random coils. Each method may have its own algorithm and database sources for prediction, enhancing our understanding of protein structure and function.

For this project, CFSSP tool and SOPMA tool were used to predict the secondary structure for the amino acid sequence for the protein.

### Tertiary structure prediction

The protein was then subjected to 3D modeling via three different web-tools viz., SWISS-MODEL (https://swissmodel.expasy.org/), Robetta (https://robetta.bakerlab.org/) and Phyre2 (http://www.sbg.bio.ic.ac.uk/phyre2/html/page.cgi?id=index) (Waterhouse et al., 2018). The modelled protein structures were downloaded in .pdb (protein data bank) format.

The quality of the predicted structure was validated by generating Ramachandran Plot for the models using the tool available in Discovery Studio Visualizer (Biovia, 2021). The Ramachandran Plot was analysed, and the best model was selected for performing docking against the ligands. The Ramachandran Z-Scores were calculated using MolProbity tool (http://molprobity.biochem.duke.edu/index.php).

The SWISS-MODEL online server utilizes information about the protein model’s geometry, interactions, and solvent potential to determine its quality. It employs the QMEAN scoring function to assess both local and overall model quality. Additionally, it provides a z-score, ranging from 0 to 1, which compares each structure’s value to the expected value (Madanagopal et al., 2022).

Phyre2 is also a modelling software which utilizes advanced remote homology detection methods for the construction of 3D models, predict probable ligand binding sites, and analyze the impact of amino acid variants, including nsSNPs, for a user provided protein sequence (Kelley et al., 2015).

The Robetta server, allows for protein structure prediction and analysis using its automated tools. Coming to structure prediction, the server firstly breaks down the submitted sequences into potential domains, which is followed by generation of structural models using either comparative modeling or *de novo* structure prediction techniques (Kim, Chivian and Baker, 2004).

### Ligand retrieval and preparation

As the title of this study suggests, two different types of ligands were used for completion of this project. The first set of ligands included the chemically modified analogues of currently used antifungals against Mucormycosis, and other Chitin Synthase inhibitors. Each synthetic drug viz., Amphotericin B, Posaconazole, Isavuconazole, Nikkomycin and Polyoxin were used as control and 10 modified analogues generated from each drug were docked against the modelled Chitin Synthase protein from *R. delemar.* The detailed chemical structure and other properties of the derivatives which were able to bind to the protein are given in the ‘Results’ section.

The 2D structures of these drug molecules were downloaded from the PubChem database and were then modified by addition/removal of different groups such as methyl, acetyl, amino, etc., using Chemsketch software, which is a molecular structure drawing application, easily downloadable from the internet (https://www.acdlabs.com/products/chemsketch/). The modified structures were downloaded in mol format and then converted to mol2 using OpenBabel (http://www.cheminfo.org/Chemistry/Cheminformatics/FormatConverter/index.html) tool. Open Babel is a software/tool that aims to simplify the conversion of chemical data between various file types and formats. This functionality is important because numerous programs have limited support for different file types or use their unique data formats. By providing a comprehensive solution, Open Babel facilitates the seamless conversion of chemical data, enabling compatibility and interoperability between diverse software applications (Madanagopal et al., 2022).

The 2D structures were then saved in 3D by using CORINA tool (https://demos.mn-am.com/corina.html). CORINA tool is an online tool which is able to generate 3D structures for small and medium sized molecules possessing drug-like properties. The modified structures were finally saved in .pdb format. The total number of modified structures were 50.

The second set of ligands were obtained from the IMPPAT database. All the ligands were downloaded in mol2 format, which is compatible with the Discovery Studio Visualizer software. The ligands were phytochemicals known for their antifungal properties (as per the details mentioned in the database) from medicinal plants such as *Acalypha indica, Azadirachta indica, Cinnamomum cassia, Datura metel,* and others (details mentioned in the table below), the IMPPAT 2.0 database located at https://cb.imsc.res.in/imppat/ was utilized. This database, known as Indian Medicinal Plants, Phytochemistry and Therapeutics 2.0 (IMPPAT 2.0), contained a comprehensive compilation of information obtained from over 100 books on traditional Indian medicine, more than 7000 published research publications, and various other sources.

Through a meticulous curation process, IMPPAT 2.0 has significantly improved and expanded upon its predecessor, IMPPAT 1.0, and currently holds the distinction of being the largest digital library focused on phytochemicals found in Indian medicinal plants. As of its version 2.0 release on June 17, 2022 (Mohanraj et al., 2018), the IMPPAT database contains data on 4010 Indian medicinal plants, 17967 phytochemicals, and 1095 therapeutic applications.

### Molecular docking and calculation of binding energies

All the prepared ligands and the prepared protein were subjected to energy minimization before molecular docking. The energy minimization was done by using the ‘Full Minimization’ tool available in Discovery Studio Visualizer.

The energy minimized ligands and proteins were then loaded for docking (LibDock) after selecting all the potential binding sites across the protein. After completion of docking, protein-ligand binding energy was calculated by selecting the ‘Calculate binding energy’ from ‘Receptor-ligand’ interactions’ in the tool bar. The more negative binding energy indicates better protein-ligand interaction. The binding energy was calculated using the equation:

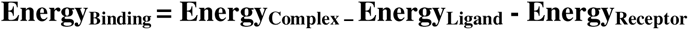

The interaction studies for top ligands were also performed after docking and binding energy calculations.

### ADMET Analysis and Toxicity testing

All the 229 ligands, which comprise of phytochemicals from IMPPAT database, commonly used anti-fungal drugs and their modified structures, were subjected to ADMET analysis using the Discovery Studio Visualizer. The following parameters were calculated for all the ligands: Human intestinal absorption after oral administration, aqueous solubility at 25°C, BBB (Blood-Brain-Barrier) penetration after absorption, CYP2D6 (Cytochrome P450 2D6) inhibition, Hepatotoxicity and Plasma-protein binding capacity. The graph was plotted between AlogP98 and the polar surface area of the molecules (Singh et al., 2021).

Subsequently, the ligand molecules were also subjected to TOPKAT profiling using Discovery Studio Visualizer. The TOPKAT wizard helped to determine the drug-likeliness of the ligands under consideration. The drug-likeliness of the ligands are determined on the basis of numerous parameters which are listed in the table 8 and 9, which shows the drug-likeliness of the selected ligands.

### Lipinski’s Rule and Veber’s rule compliance

Lipinski’s rule of five, which is also known as Pfizer’s rule of five, or more commonly known as the rule of five (RO5) is taken as a thumb rule for evaluating the drug-likeliness property of a ligand, which in turn determines if a particular chemical compound has the ability to be effective within humans after oral consumption. This rule was given by Christopher Lipinski in late 90’s which is based on the observation that most of the drugs which are orally administered are relatively small and are moderately lipophilic in nature.

The following five conditions are specified by Lipinski for a drug to be orally bioavailable:

- Number of Hydrogen-bond donors is not more than 5.
- Number of Hydrogen-bond acceptors is not more than 10.
- Molecular weight is not more than 500.
- LogP is not more than 5.

Along with Lipinski’s rule, Veber’s rule is another criterion which needs to be satisfied for a drug molecule

- The number of total rotatable bonds is not more than 10.
- The polar surface area is not more than 140Å^2^ or
- The sum of Hydrogen-bond donors and acceptors is not more than 15.

The docked ligands were tested for their drug-likeliness based on these 2 parameters, i.e., both Lipinski’s rule and Veber’s rule compliance.

## RESULTS AND DISCUSSION

### Protein sequence retrieval

The protein sequence of interest, i.e., Chitin synthase of *R. delemar* was derived from NCBI in FASTA format (GenBank ID: EIE85441.1).

### Analysis of Physicochemical properties of the protein

The ExPasy’s ProtParam server was used to calculate the various physico-chemical parameters for the amino acid sequence coding for the Chitin Synthase enzyme of *Rhizopus delemar.* The following characteristics were determined for the protein (table 2):

**Table 1.**
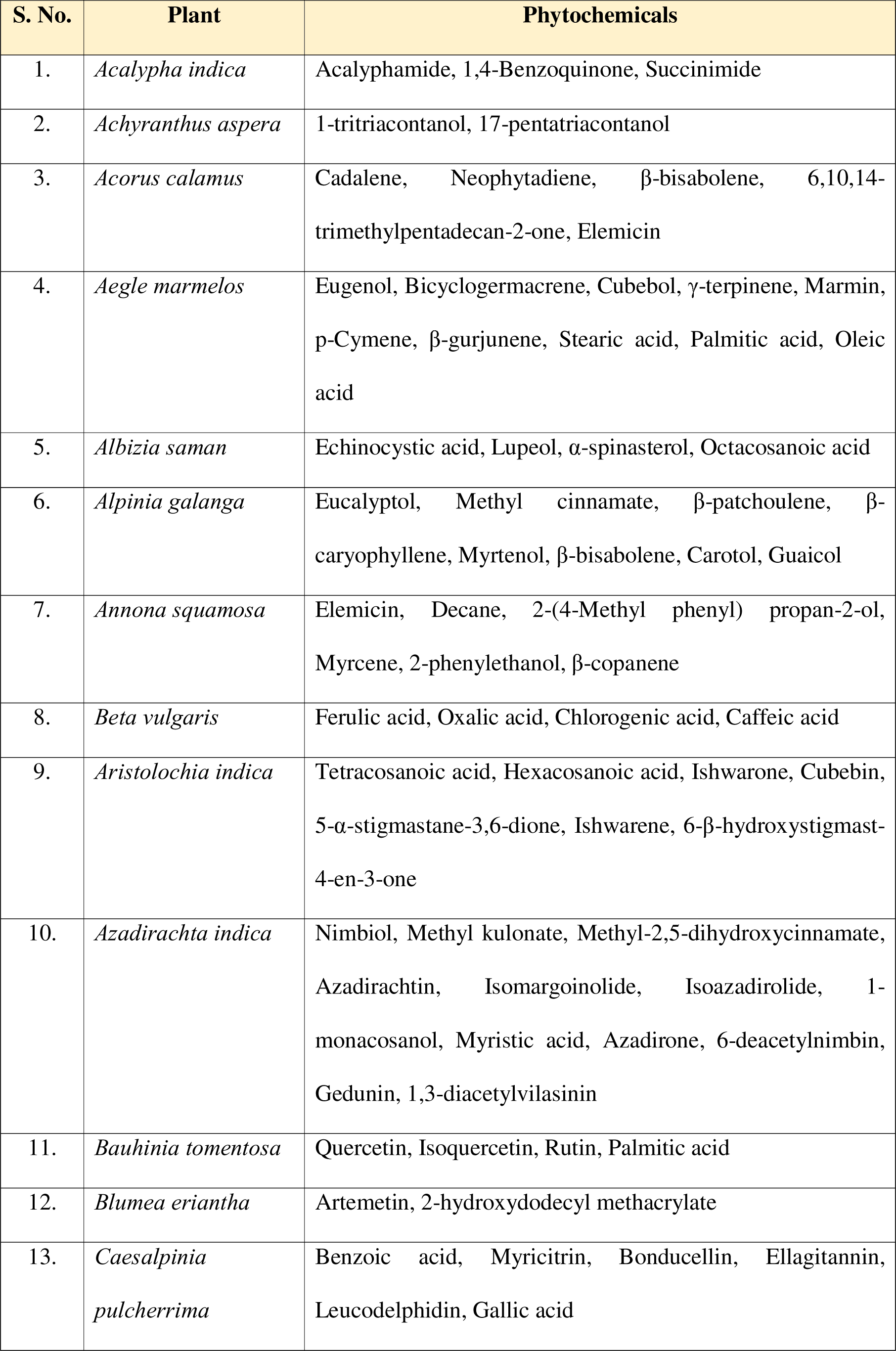
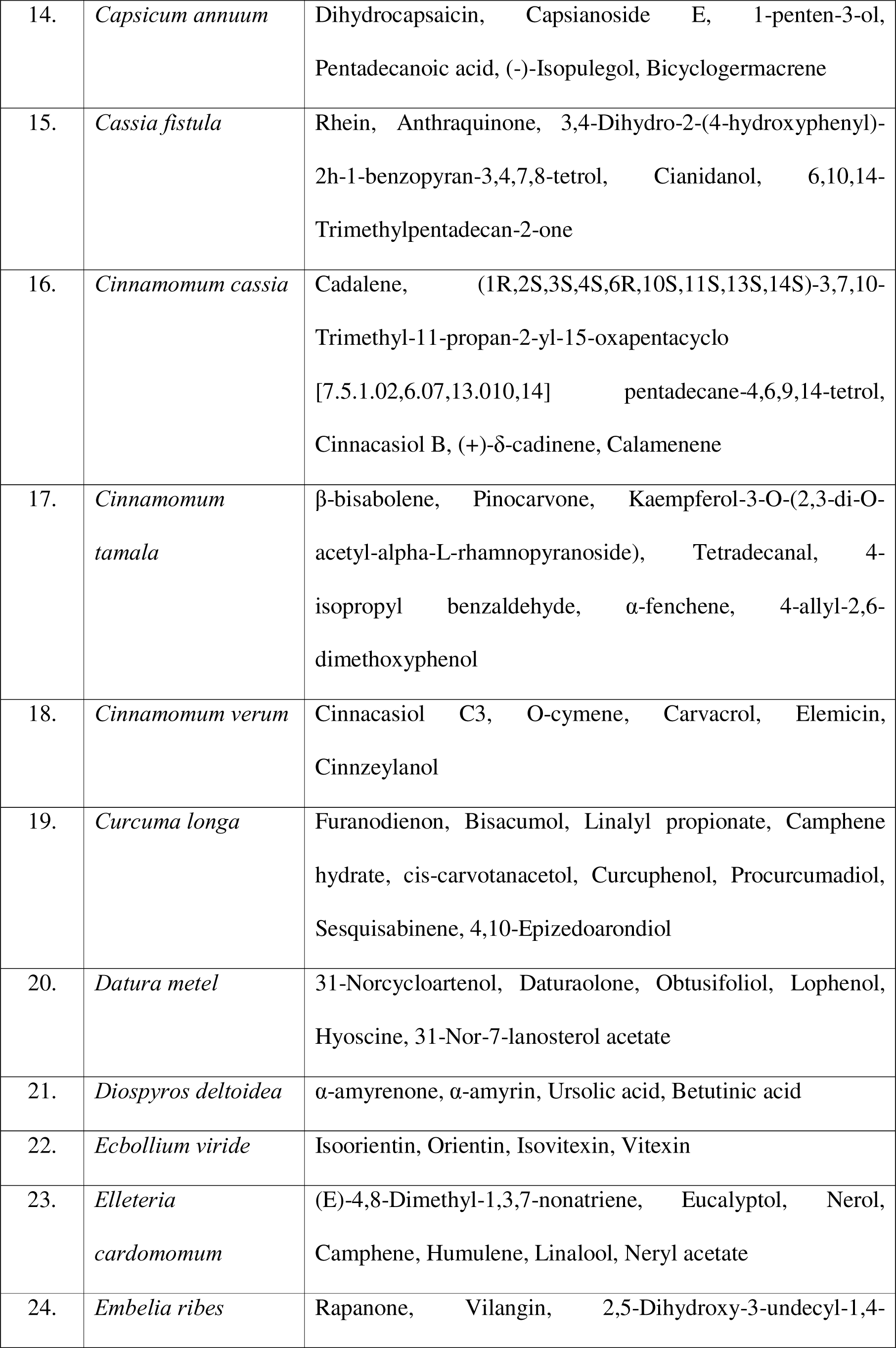
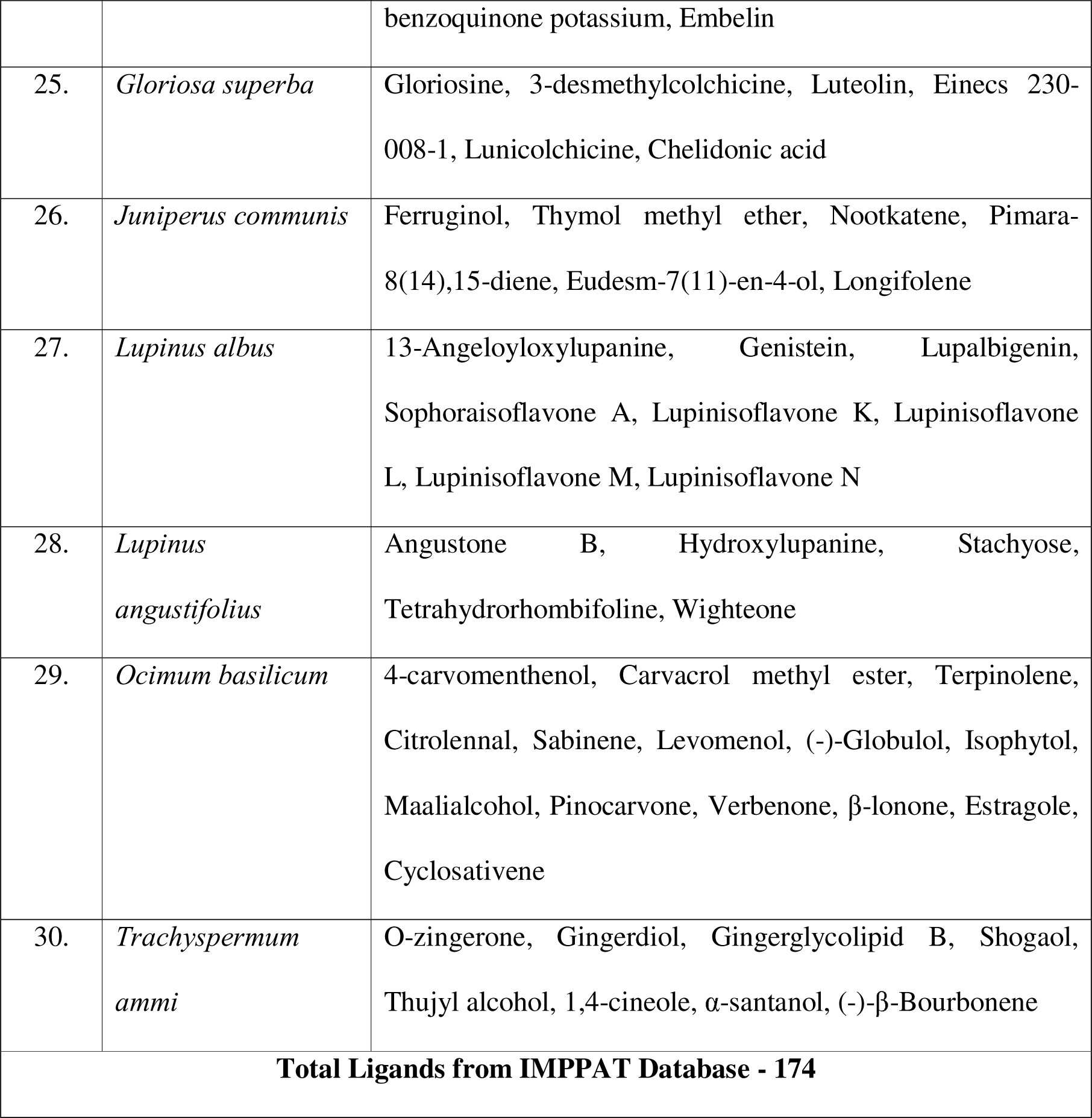
Phytochemicals from IMPPAT Database with antifungal properties obtained from 30 different plants.

**Table 2.**
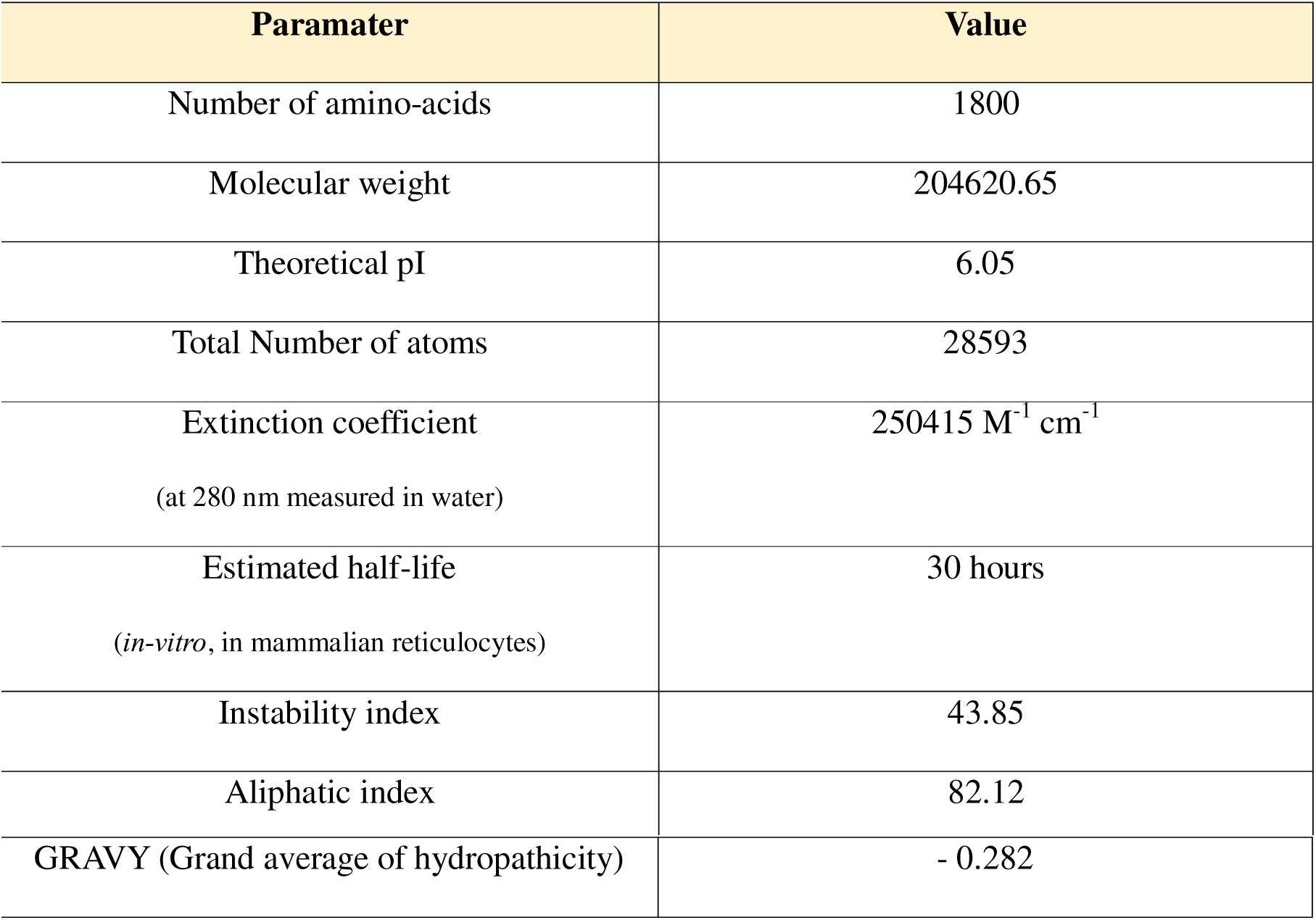
Physico-chemical characteristics of the protein as calculated by ProtParam server.

### Secondary Structure Prediction

The secondary structure for the protein was predicted using both the CFSSP tool and SOPMA tool available on the internet. CFSSP suffers from the handicap that only proteins up to 1000 amino acids can be used for analysis, thereby limiting its use for longer sequences. On the other hand, it was observed that SOPMA allows even longer proteins to get their secondary structure predicted at the expense of time. The results obtained suggested that the protein was predominantly having Random coils in its structure which accounted for approximately 44.1% of the protein, followed by α-helices which accounted for almost 39.17% of the protein. The detailed analysis is given in the figure 3.

**Fig. 1.**
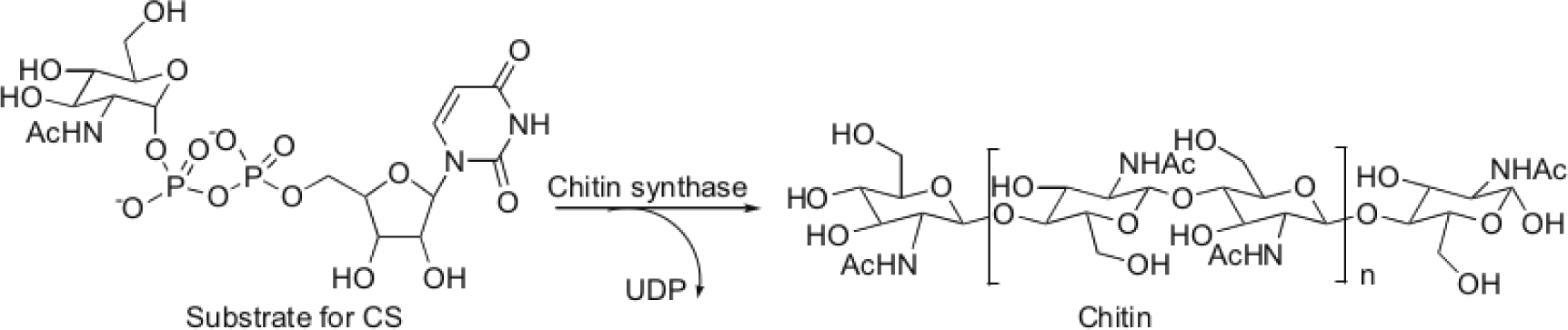
Mechanism of Chitin Synthesis (Chaudhary, Tupe and Deshpande, 2013)

**Fig. 2.**
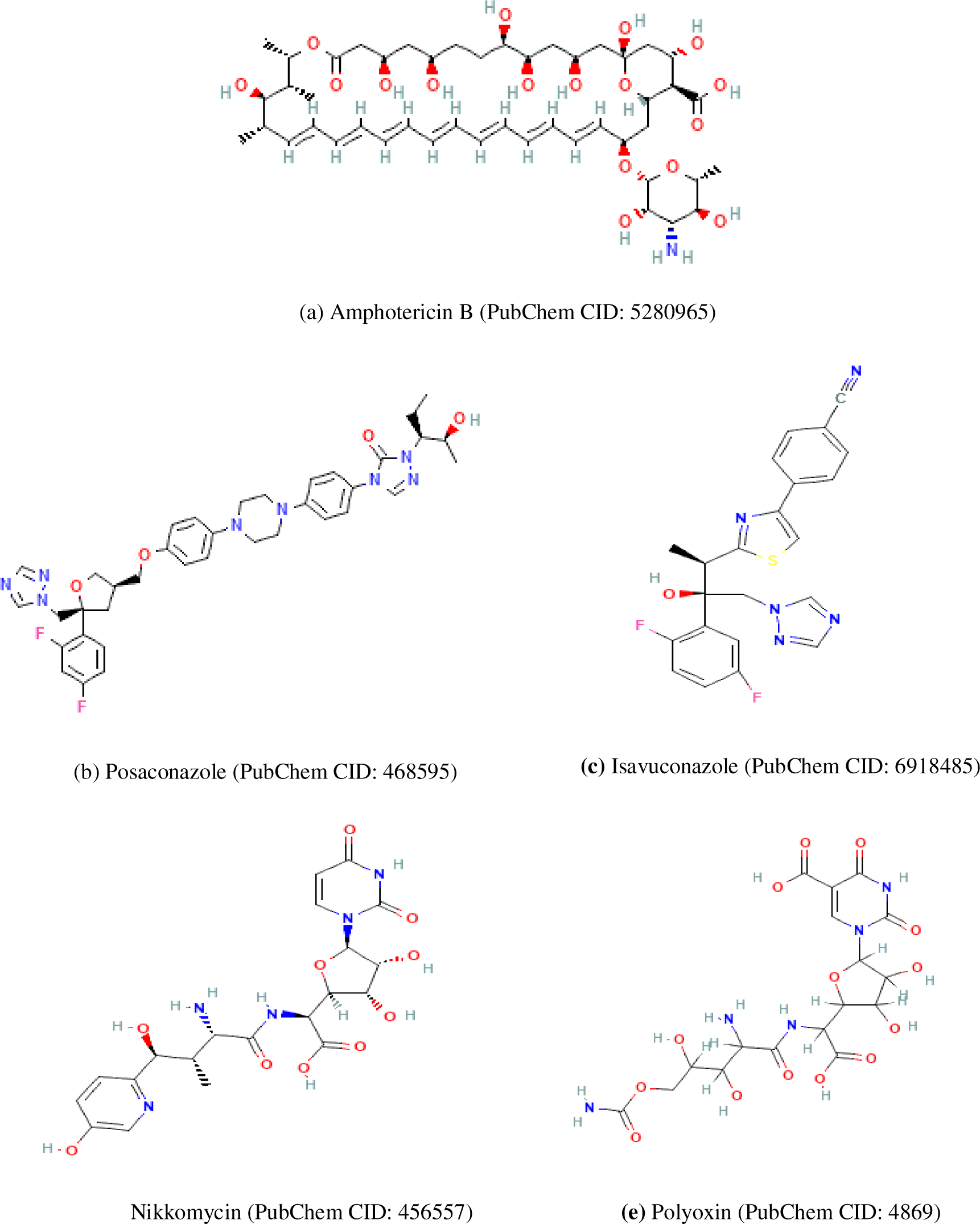
**(a) to 2(c)** Commonly used medications against Mucormycosis; **(d) and (e)** Known Chitin Synthase inhibitors Source: PubChem (https://pubchem.ncbi.nlm.nih.gov/) (Kim et al., 2023)

**Fig. 3.**
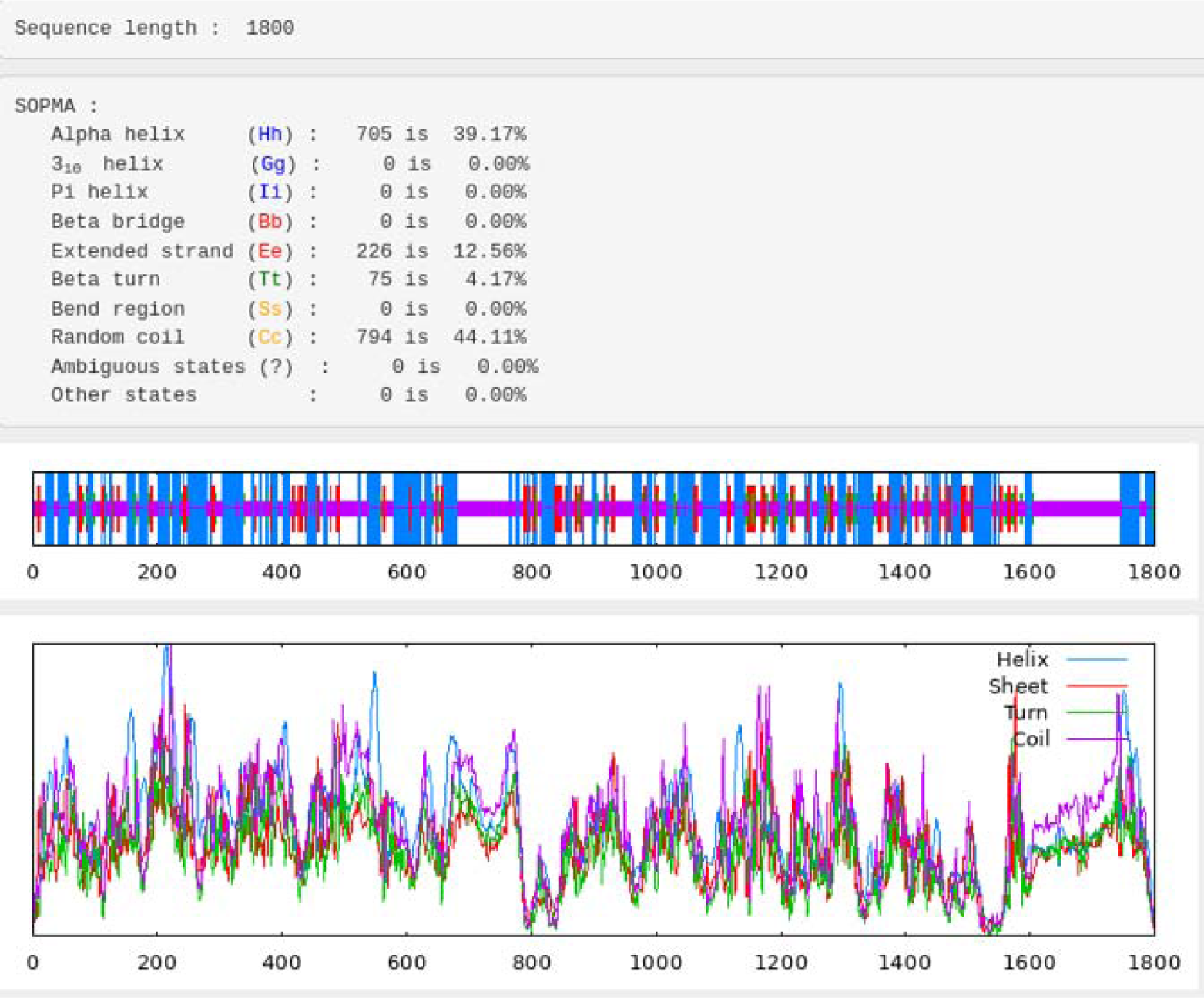
Secondary structure prediction results for the protein using SOPMA

### Tertiary structure prediction

The tertiary structure for the protein (Chitin Synthase) was modelled using different tools available in the internet *viz.,* SWISS-MODEL, Phyre2 and Robetta. A total of 3 different models were selected, one from each tool, out of the number of models generated by each tool. The generated models were saved in pdb format and were validated using Ramachandran Plot and Ramachandran Z-score values (generated using MolProbity tool). The Ramachandran plots were generated using the Ramachandran plot generation tool in Discovery Studio Visualizer. The results are given in table 3.

**Table 3.**
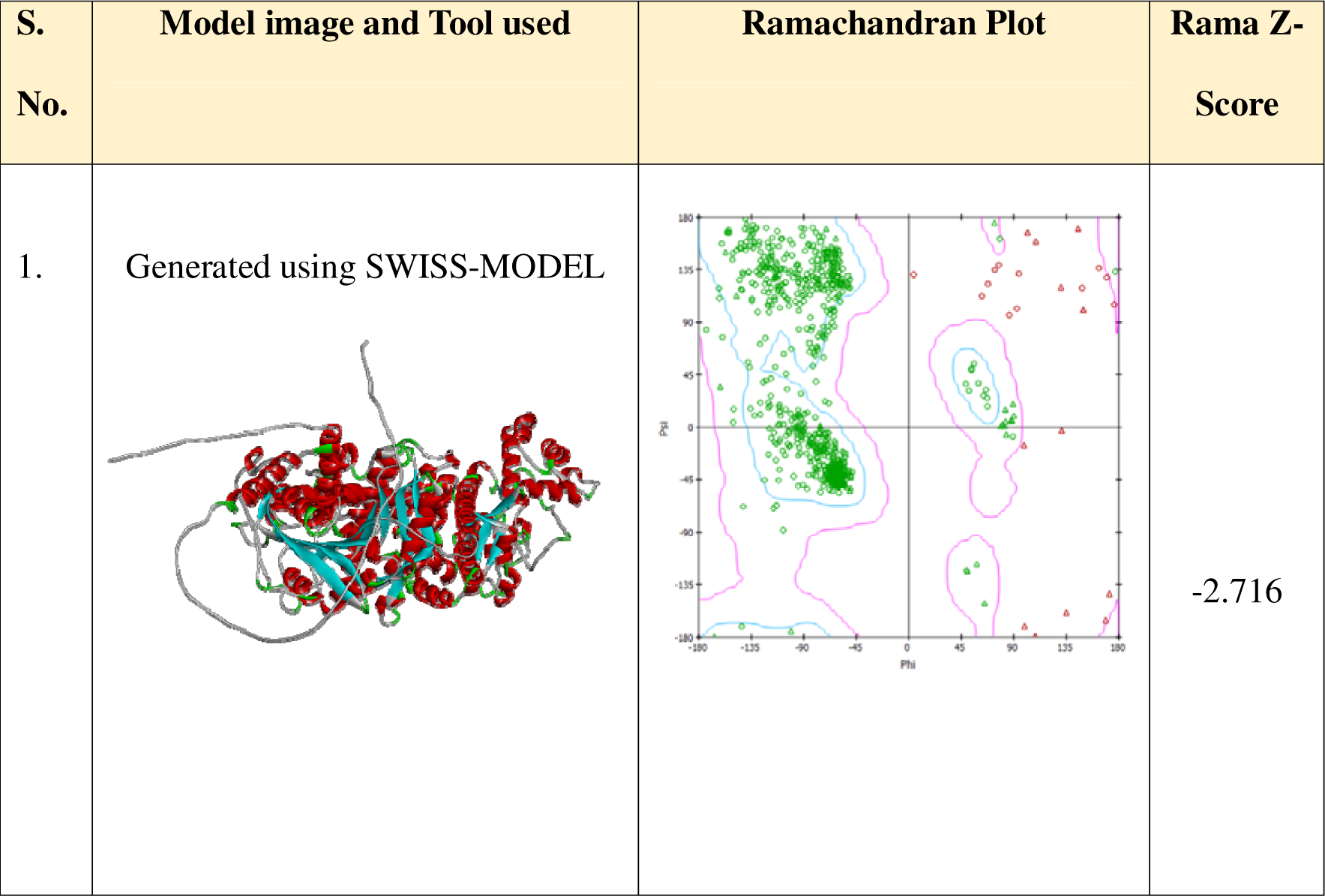
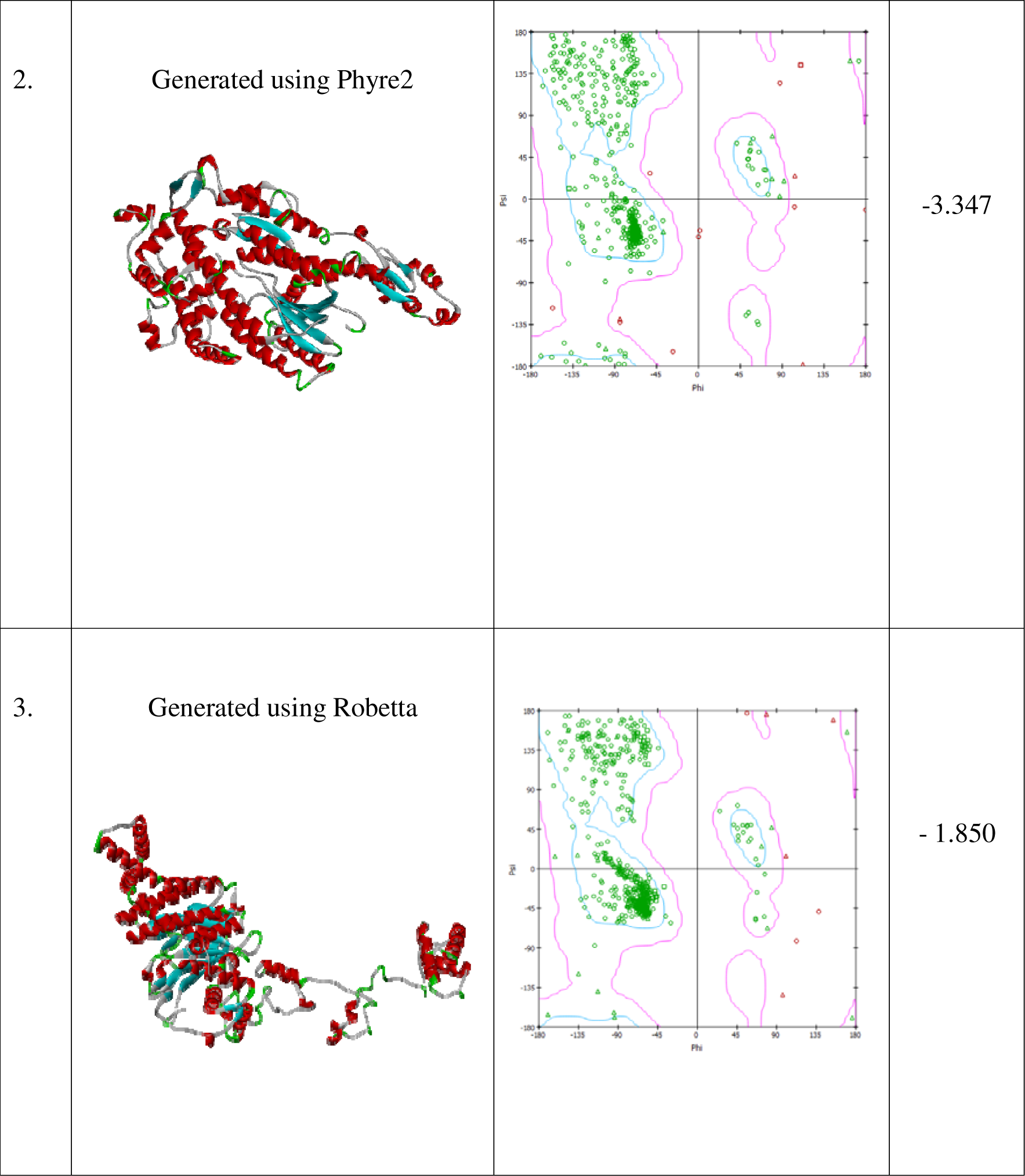
3D structures for the protein generated using different tools and their validation using Ramachandran Plot.

| S. No. | Model image and Tool used | Ramachandran Plot | Rama Z-Score |
| --- | --- | --- | --- |
| 1. | Generated using SWISS-MODEL |  | -2.716 |
| 2. | Generated using Phyre2 |  | -3.347 |
| 3. | Generated using Robetta |  | - 1.850 |

Considering all the other factors in addition to Ramachandran Plot and Rama Z-score, such as % coverage, sequence identity and sequence similarity and homology, the model generated by using SWISS-MODEL online tool was adjudged to be the best model, and was used for the docking studies.

### Docking results of anti-fungal drugs and their derivatives (modified drugs)

A total of 55 drugs (5 anti-fungal drugs, and their modified analogues (10 derivatives for each drug) were docked against the modelled protein using the LibDock tool of Discovery studio visualizer. From these 55 drugs, only 6 were docked to the protein and 49 of them failed to dock, indicating their failure to bind to the protein. Out of these 6 docked drugs, 1^st^ modified derivative of Posaconazole (Posaconazole1) with a LibDock score of 107.862 was the top binding ligand followed by 1^st^ modified derivative of Polyoxin (Polyoxin1), with a LibDock score of 105.194. Fig. 4(a) to (f) below shows the structure of the docked drugs, in descending order of their LibDock score.

**Fig. 4.**
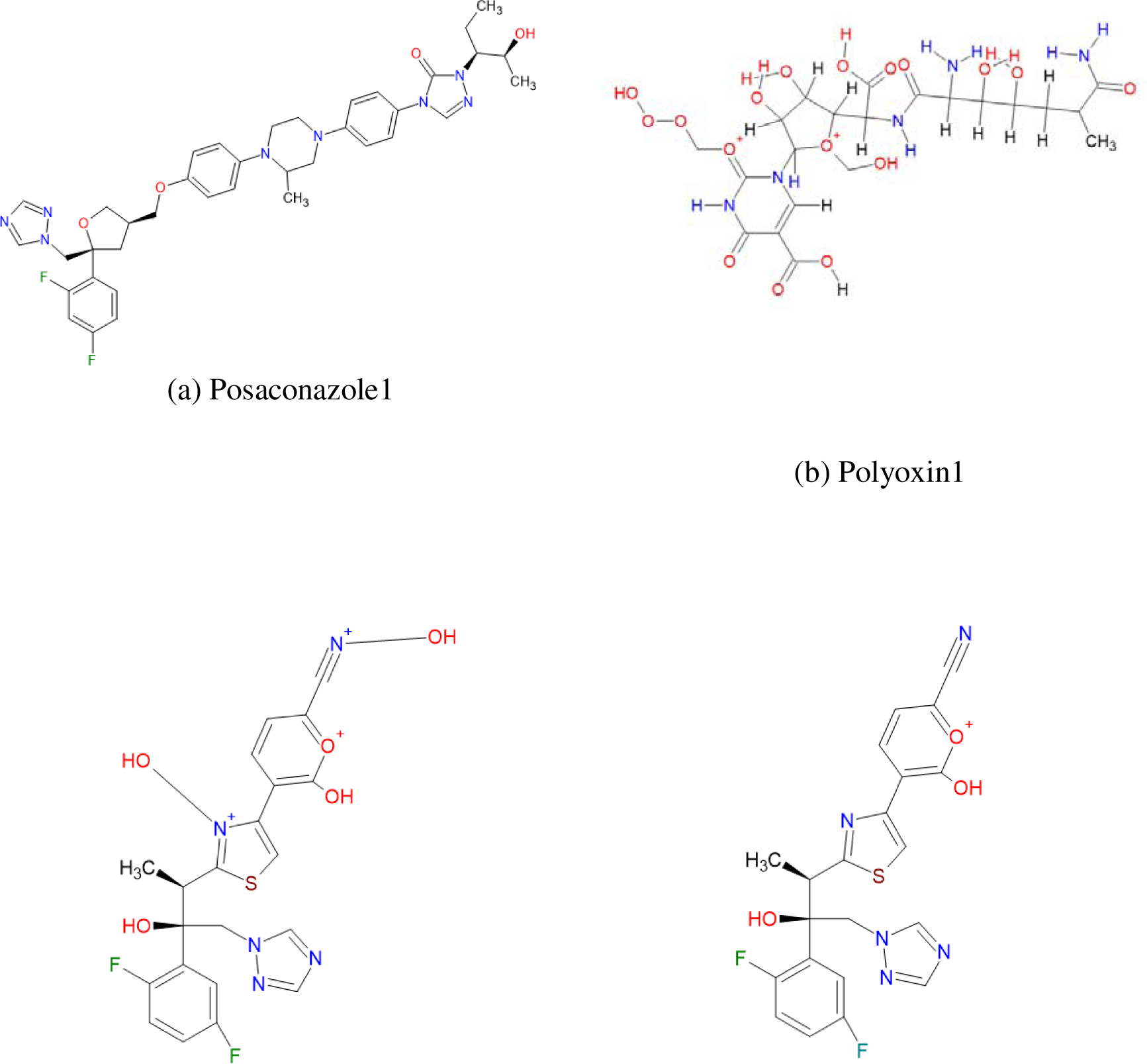
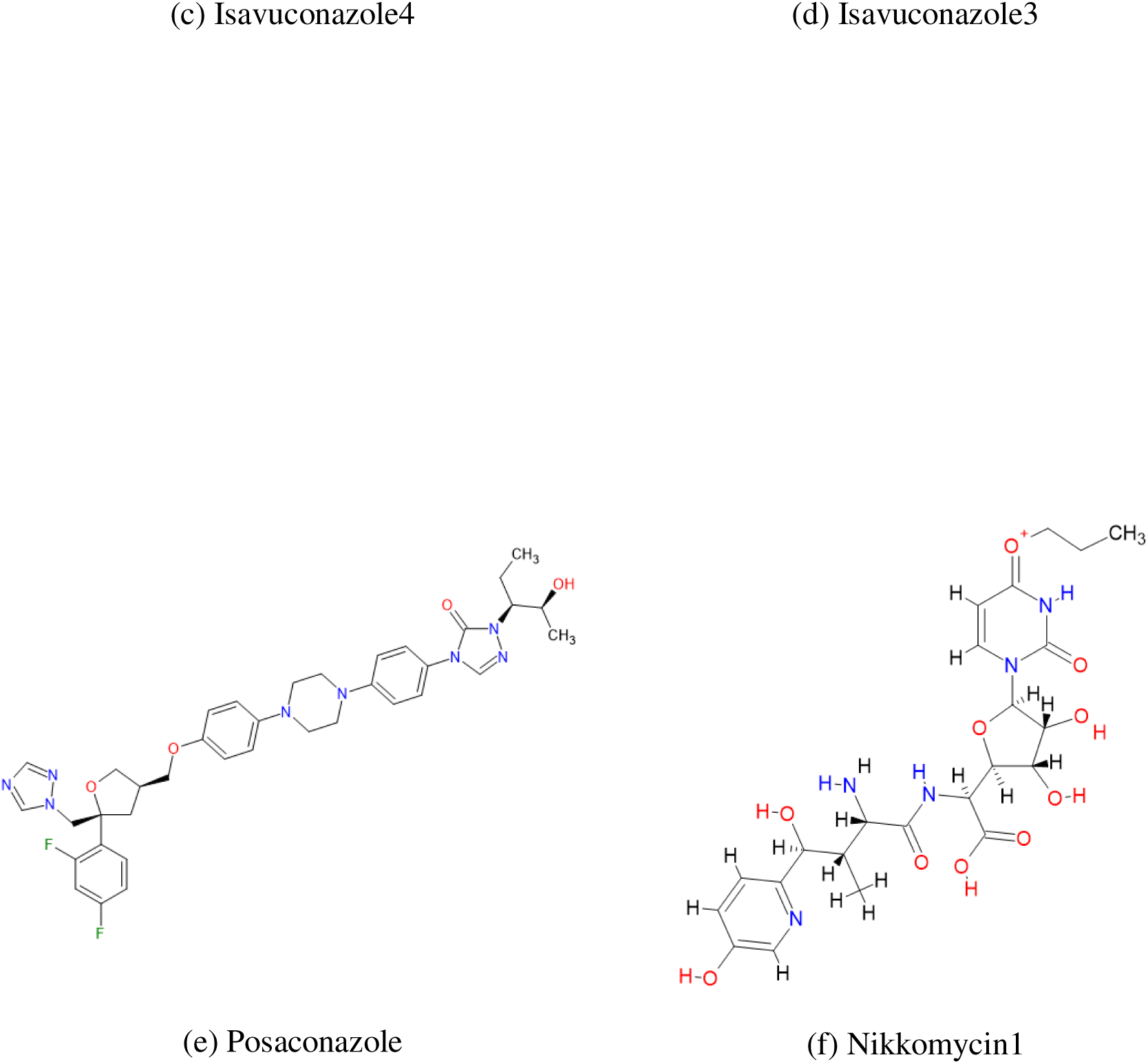
**(a) to (f)** Drugs/Modified drugs which were able to bind to the protein

Table 4 below shows the molecular descriptors of the above-mentioned drugs. Details such as Molecular formula, Molecular weight, Molecular Composition, Number of atoms, etc. are recorded.

**Table 4.**
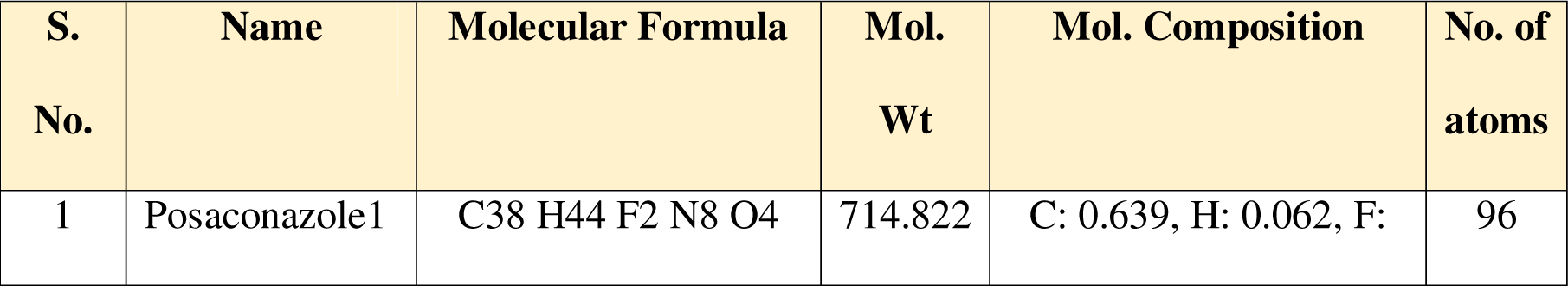
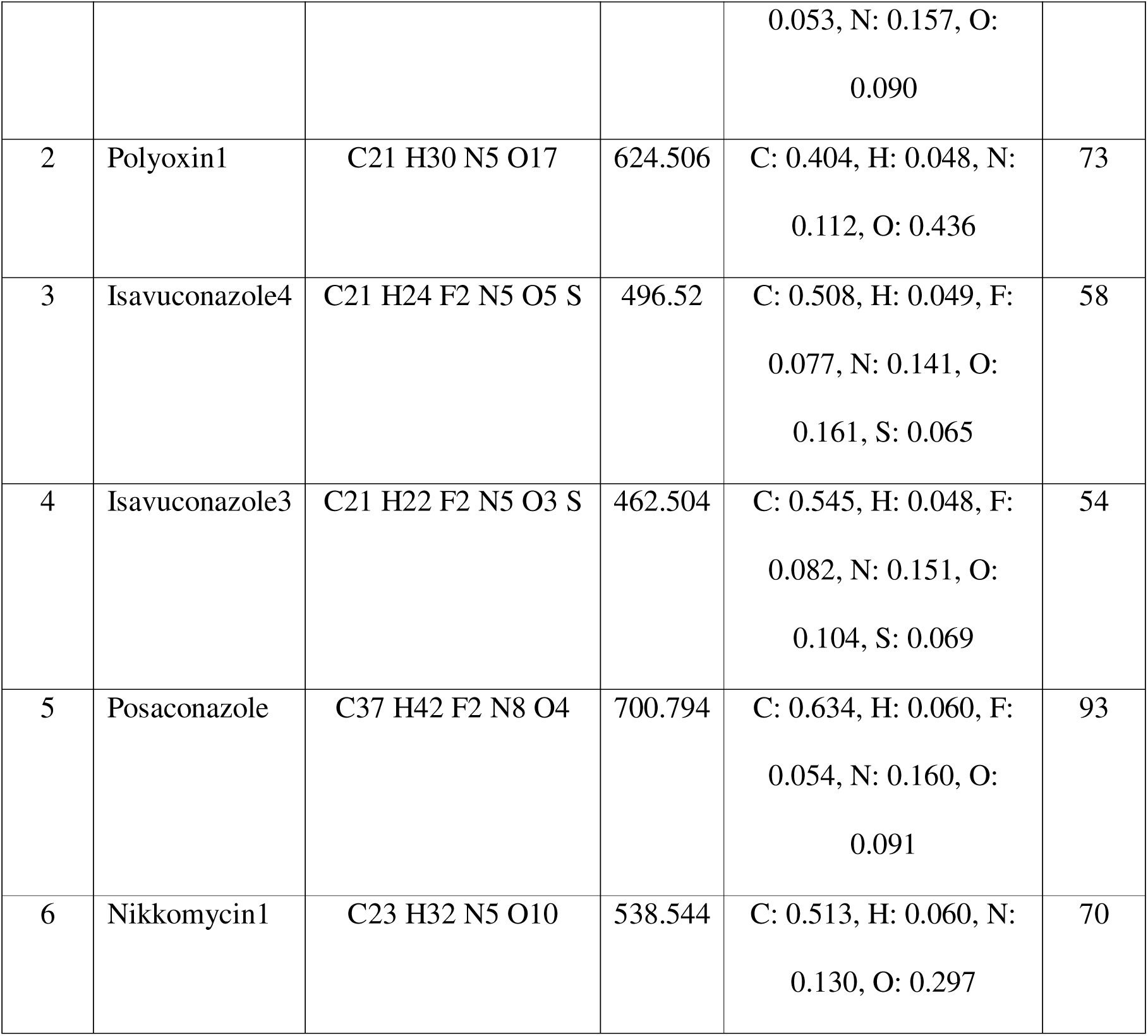
Molecular descriptors of anti-fungal drug/drug derivatives.

The LibDock score and the binding energies for each of the above drugs docked are given in table 5. It is to be noted that only top pose of drugs/drug derivatives were taken for our studies, as they have the highest LibDock scores in comparison to other poses. Posaconazole had the highest LibDock score among all 6 drug/drug derivatives.

**Table 5.** Binding affinity and Binding energy of anti-fungal drugs/drug derivatives.

| S.<br>No. | Name | LibDock<br>Score of<br>top pose | Binding<br>Energy | Ligand<br>Energy | Protein<br>Energy | Complex<br>Energy |
| --- | --- | --- | --- | --- | --- | --- |
| 1 | Posaconazole1 | 107.862 | 995.922 | 158.356 | -57347.6 | -56193.3 |
| 2 | Polyoxin1 | 105.194 | 19537.2 | 462.253 | -57347.6 | -37348.1 |
| 3 | Isavuconazole4 | 90.2841 | 1825.23 | 899.916 | -57347.6 | -54622.4 |
| 4 | Isavuconazole3 | 87.6533 | 4006.8 | 133.129 | -57347.6 | -53207.6 |
| 5 | Posaconazole | 85.7407 | 581.866 | 246.291 | -57347.6 | -56519.4 |
| 6 | Nikkomycin1 | 76.1422 | 12281.7 | 67.4276 | -57347.6 | -44998.4 |

### Docking results of phytochemicals

A total of 174 anti-fungal compounds from 30 different plants were docked against the prepared protein. Only 19 out of 174 were docked to the protein and 155 failed to dock. Out of these 19 docked phytochemicals, 1-Nonacosanol had the highest LibDock score of 118.483, followed by Cubebin, with a LibDock score of 104.206. Fig. 5(a) to (s) below shows the structure of the docked drugs, in descending order of their LibDock score.

**Fig. 5.**
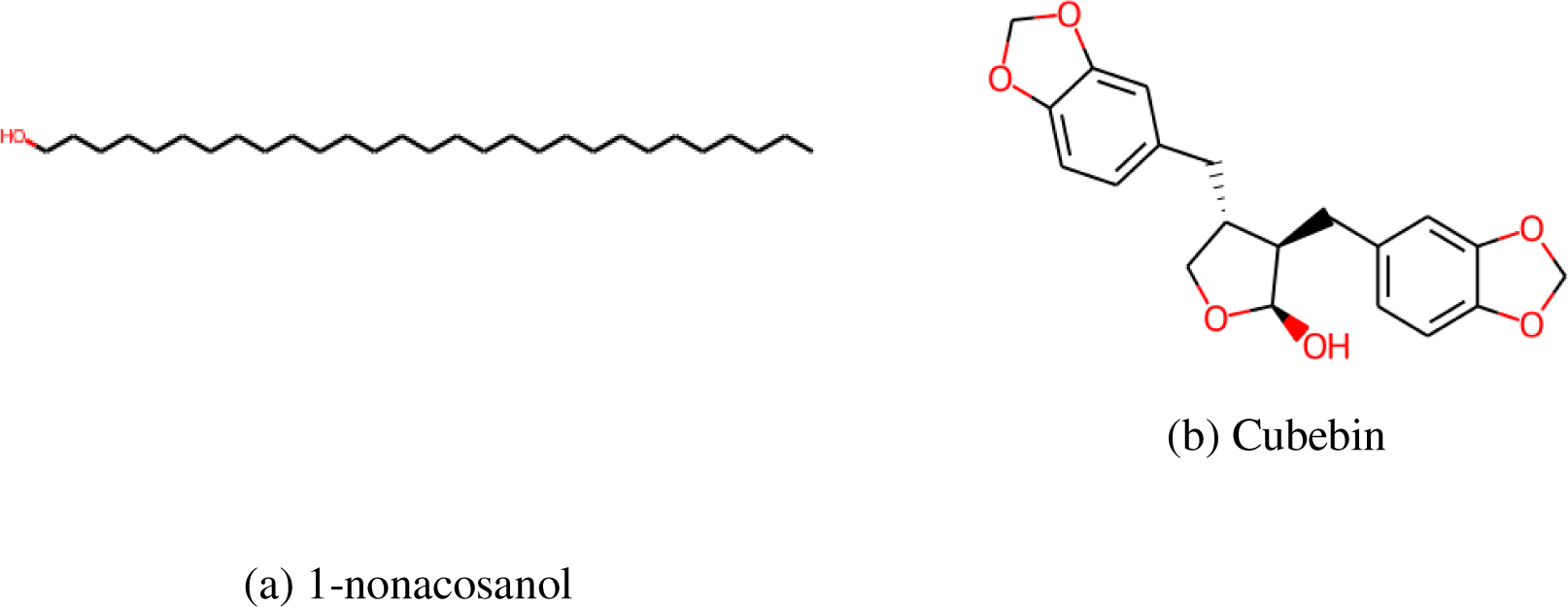
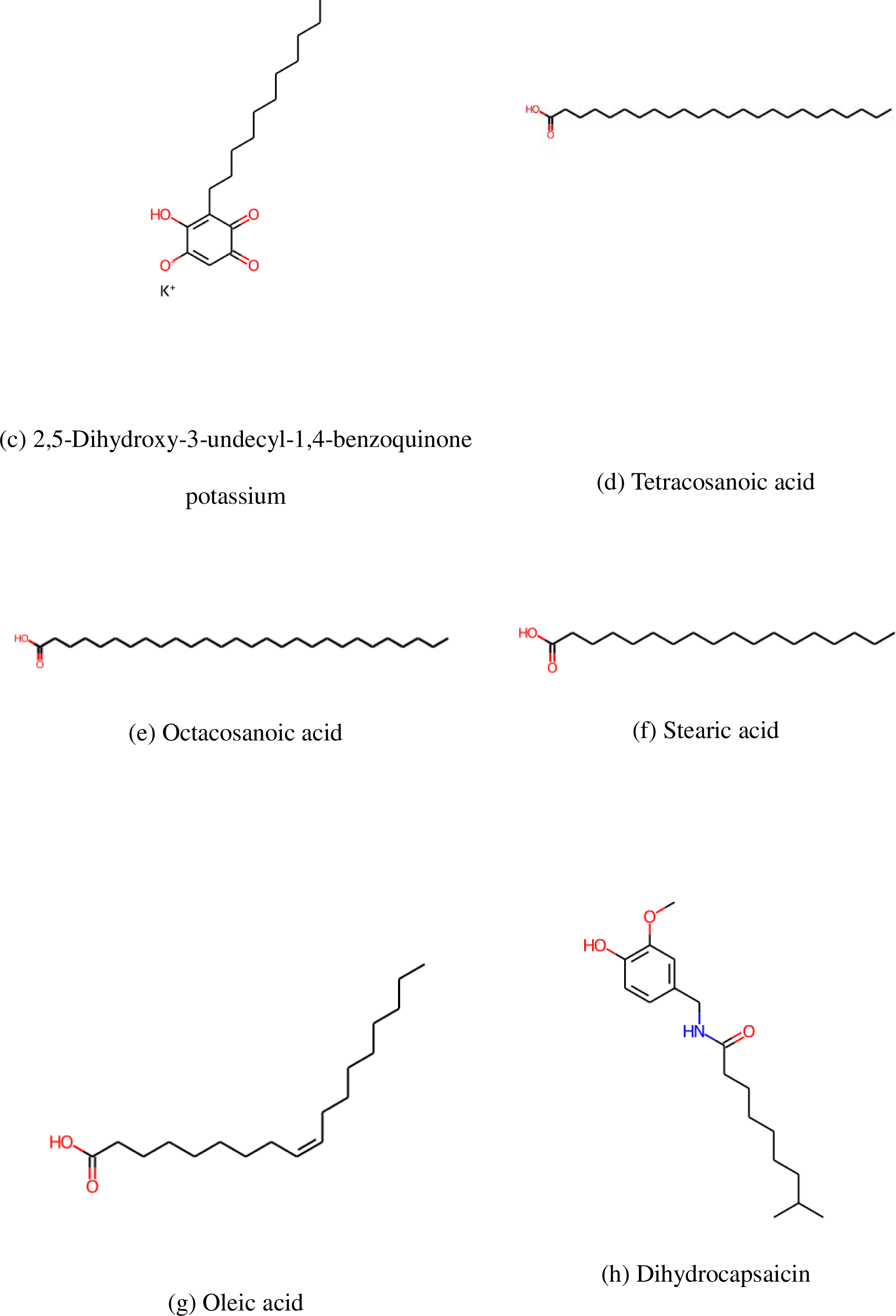
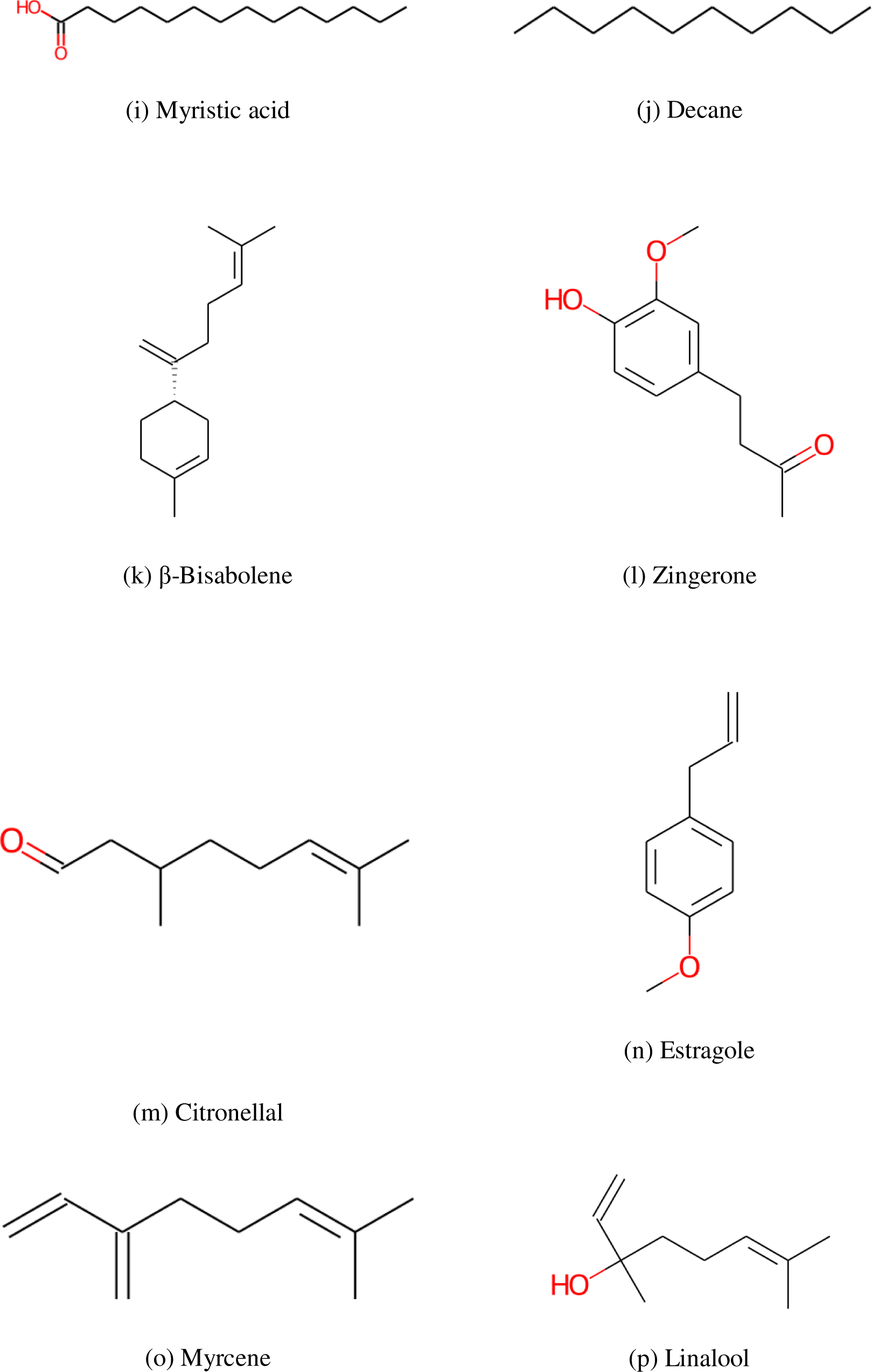
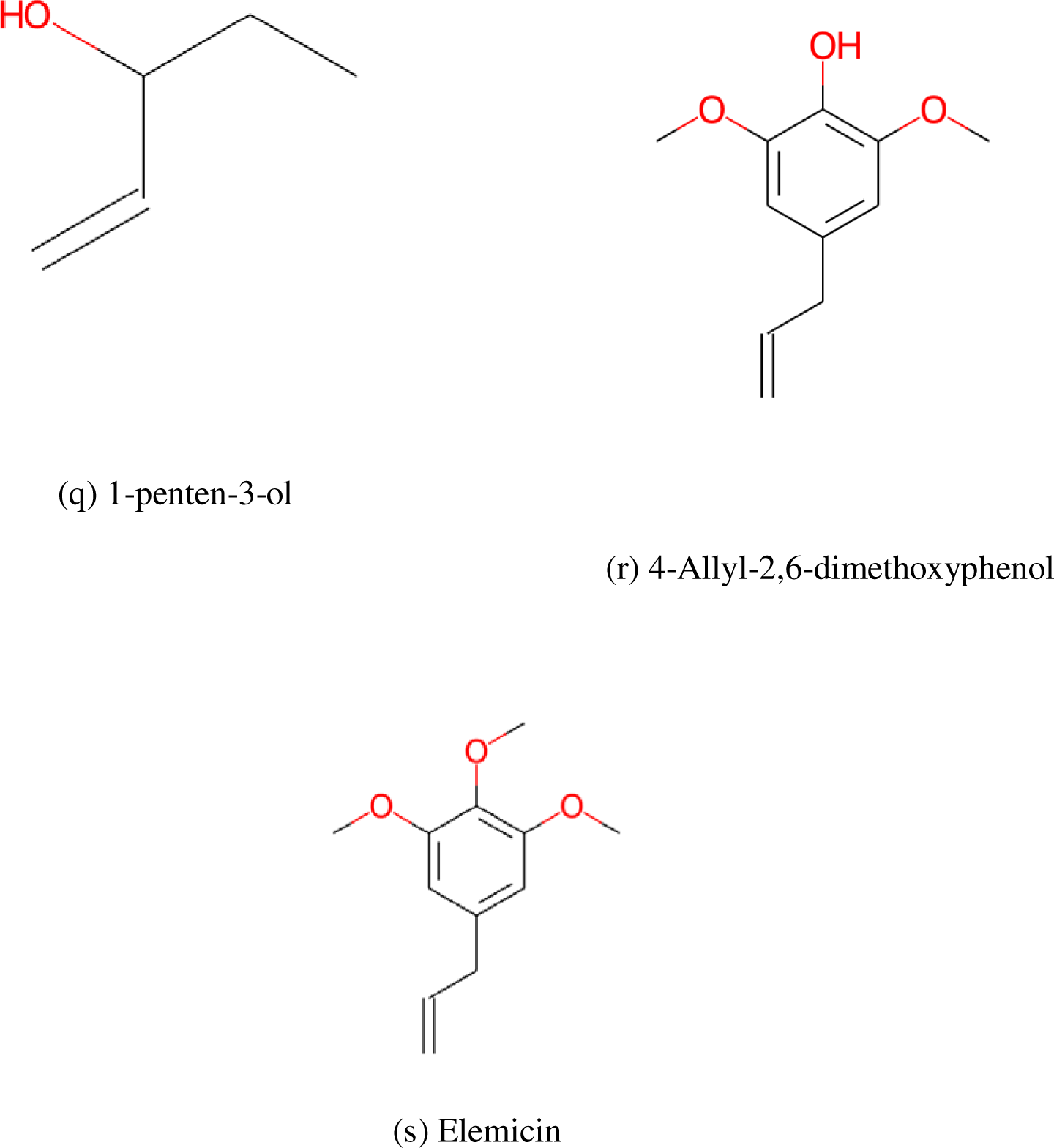
**(a) to (s)** Structure of the phytochemicals which were able to bind to the protein

Table 6 shows the molecular descriptors of the above-mentioned phytochemicals. Details such as Molecular formula, Molecular weight, Molecular Composition, Number of atoms, etc. are recorded.

**Table 6.**
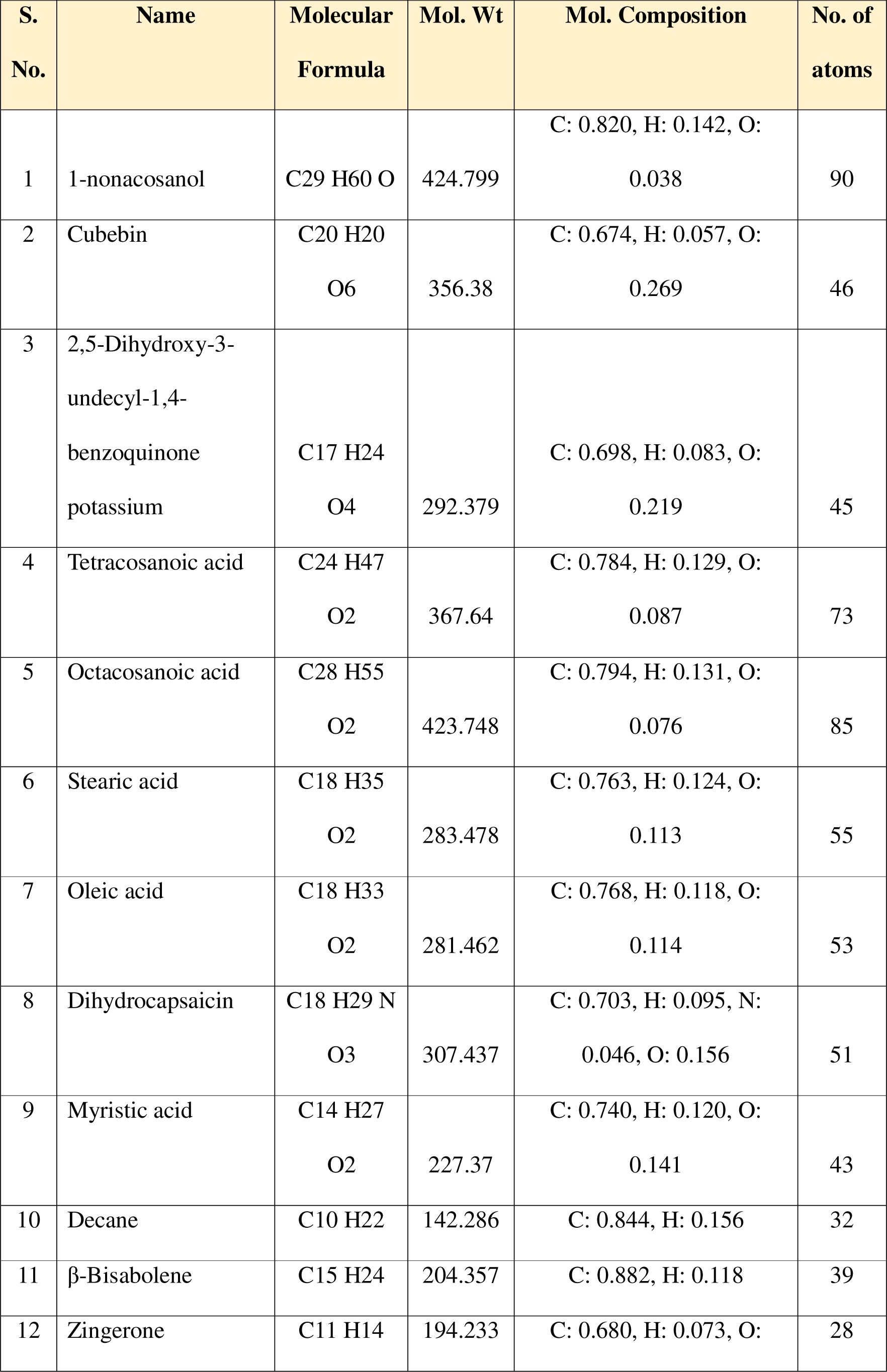
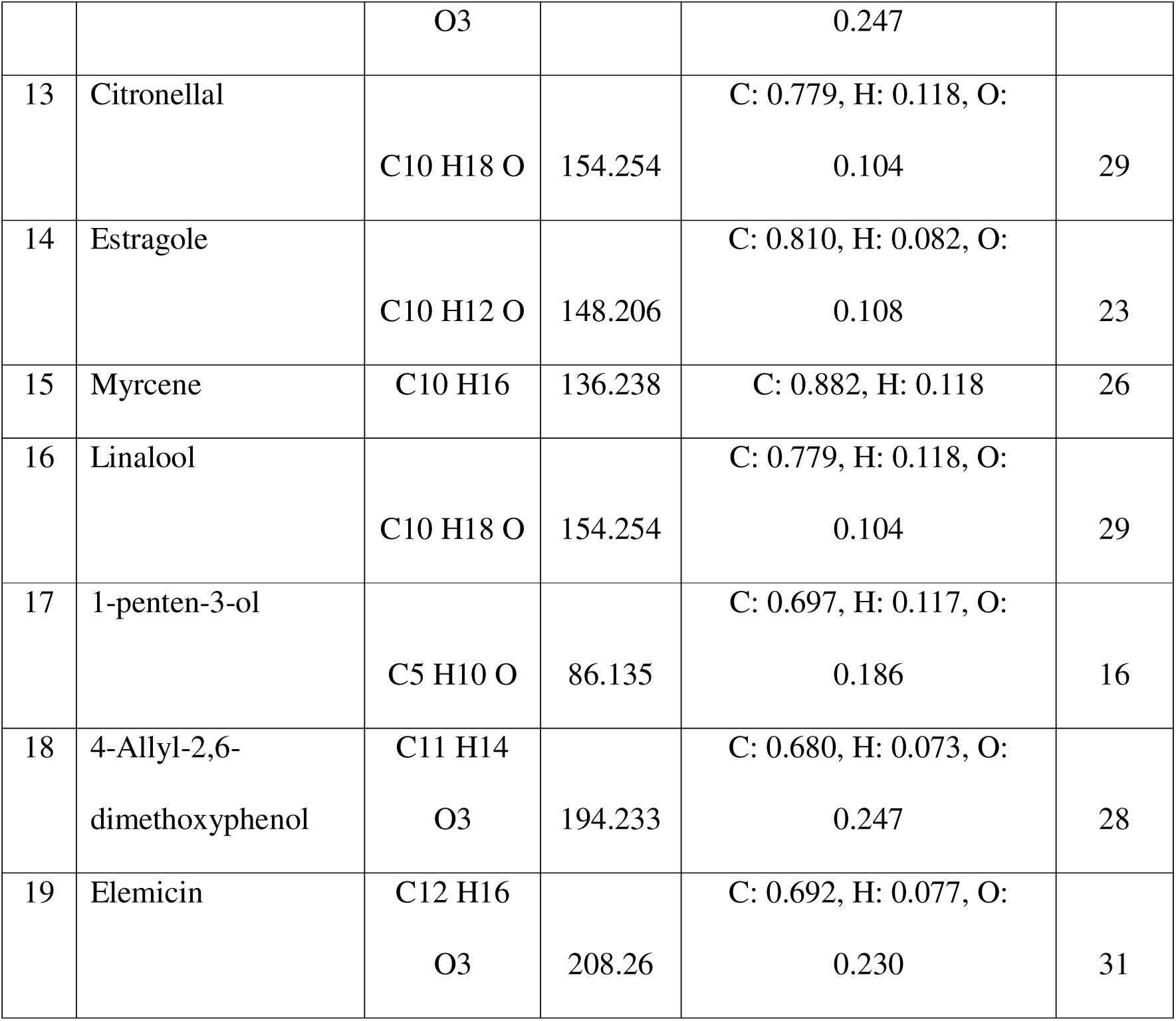
Molecular descriptors of anti-fungal phytochemicals.

The LibDock score and the binding energies for each of the above phytochemicals docked are given in table 7. It is to be noted that only top pose of drugs/drug derivatives were taken for our studies, as they have the highest LibDock scores in comparison to other poses. 1-nonacosanol had the highest LibDock score among all 19 phytochemicals.

**Table 7.** Binding affinity and Binding energies of anti-fungal phytochemicals.

| S. No. | Name | LibDock Score of top pose | Binding Energy | Ligand Energy | Protein Energy | Complex Energy |
| --- | --- | --- | --- | --- | --- | --- |
| 1 | 1-nonacosanol | 118.483 | 2872.15 | 21.9278 | -57347.6 | -54453.5 |
| 2 | Cubebin | 104.206 | 77.5743 | 103.33 | -57347.6 | -57166.7 |
| 3 | 2,5-Dihydroxy-3-undecyl-1,4-benzoquinone potassium | 103.888 | 23943.9 | 153.722 | -57347.6 | -33249.9 |
| 4 | Tetracosanoic acid | 101.225 | 242751 | -0.0157 | -57347.6 | 185403 |
| 5 | Octacosanoic acid | 96.3122 | 3328.96 | 28.0943 | -57347.6 | -53990.5 |
| 6 | Stearic acid | 93.0352 | 882.477 | 33.8405 | -57347.6 | -56431.3 |
| 7 | Oleic acid | 90.0019 | 11723.4 | 35.5725 | -57347.6 | -45588.6 |
| 8 | Dihydrocapsaicin | 84.4284 | 1076.47 | 12.8054 | -57347.6 | -56258.3 |
| 9 | Myristic acid | 78.2806 | 338.252 | 21.8824 | -57347.6 | -56987.4 |
| 10 | Decane | 76.0367 | 1794.33 | 5.6698 | -57347.6 | -55547.6 |
| 11 | β-Bisabolene | 75.4589 | 50118.4 | 117.199 | -57347.6 | -7112 |
| 12 | Zingerone | 69.1929 | 422.621 | 7.8195 | -57347.6 | -56917.1 |
| 13 | Citronellal | 66.8427 | 17654.6 | 60.0491 | -57347.6 | -39633 |
| 14 | Estragole | 65.5456 | 192.678 | 16.0557 | -57347.6 | -57138.8 |
| 15 | Myrcene | 57.5669 | 153.873 | 62.3028 | -57347.6 | -57131.4 |
| 16 | Linalool | 54.9492 | 2412.64 | 64.1426 | -57347.6 | -54870.8 |
| 17 | 1-penten-3-ol | 48.1763 | 4941.79 | 22.5331 | -57347.6 | -52383.3 |
| 18 | 4-Allyl-2,6-dimethoxyphenol | 45.0608 | -10.667 | 388.467 | -57347.6 | -56969.8 |
| 19 | Elemicin | 36.2331 | 3784.52 | 162.232 | -57347.6 | -53400.8 |

**Table 8.**
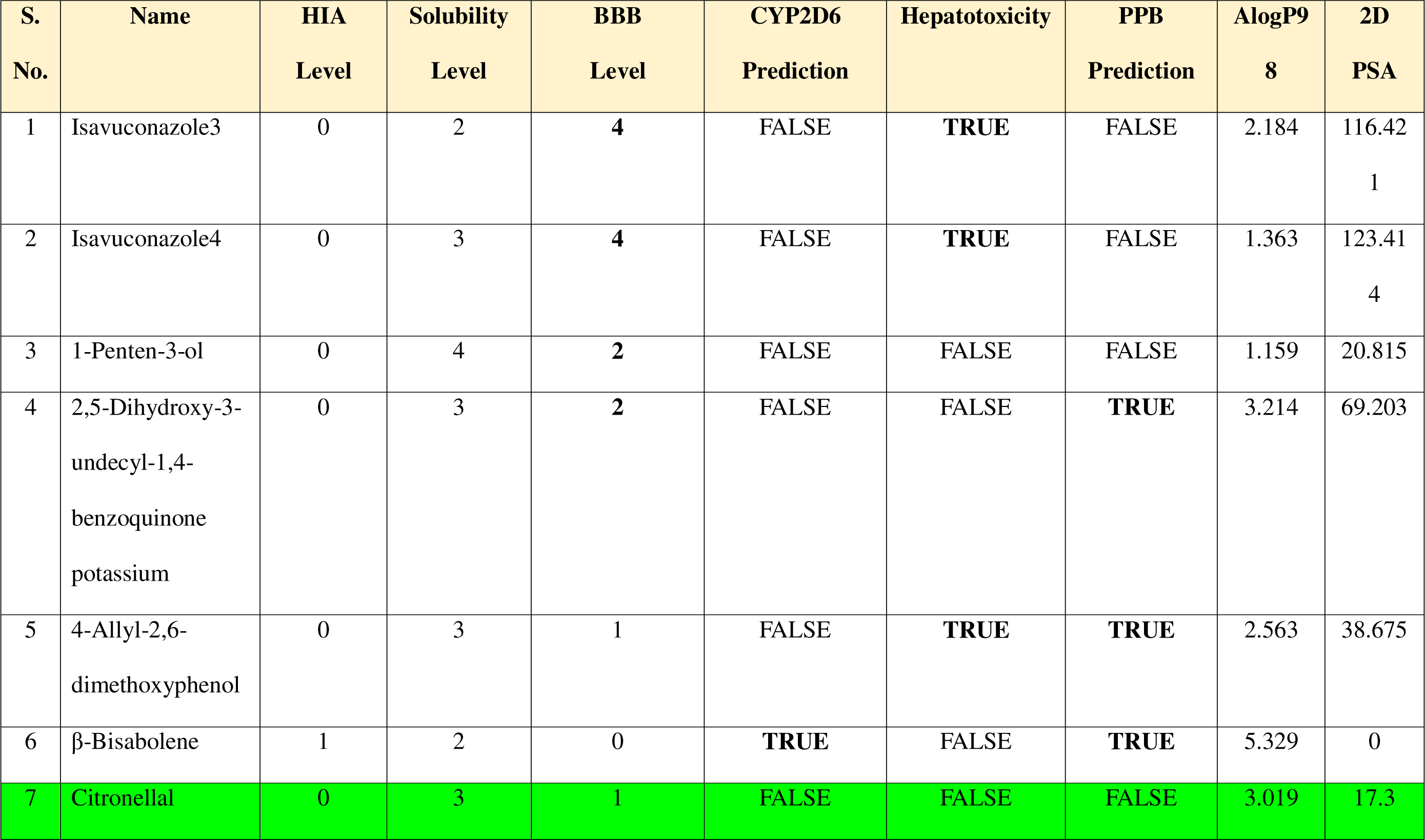
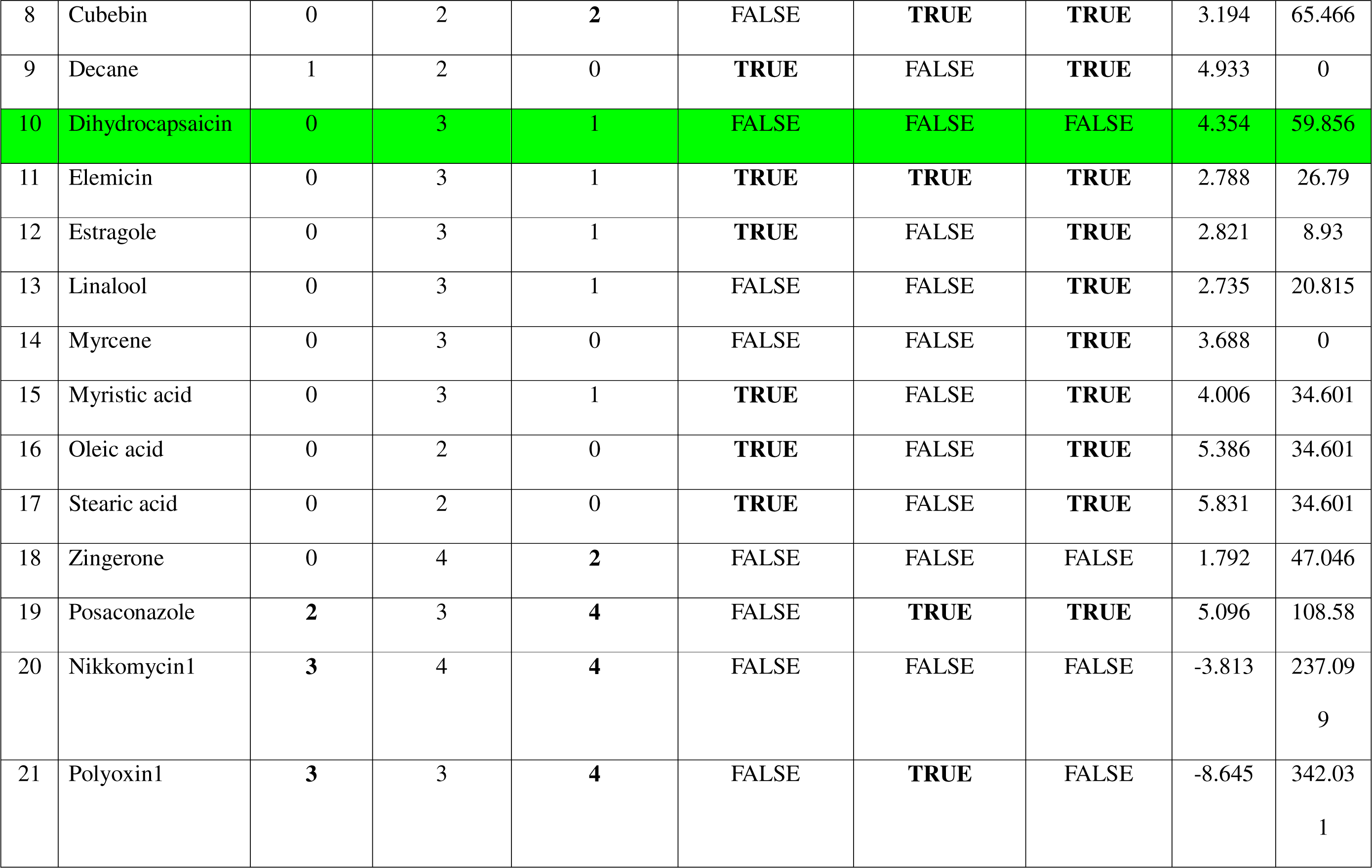
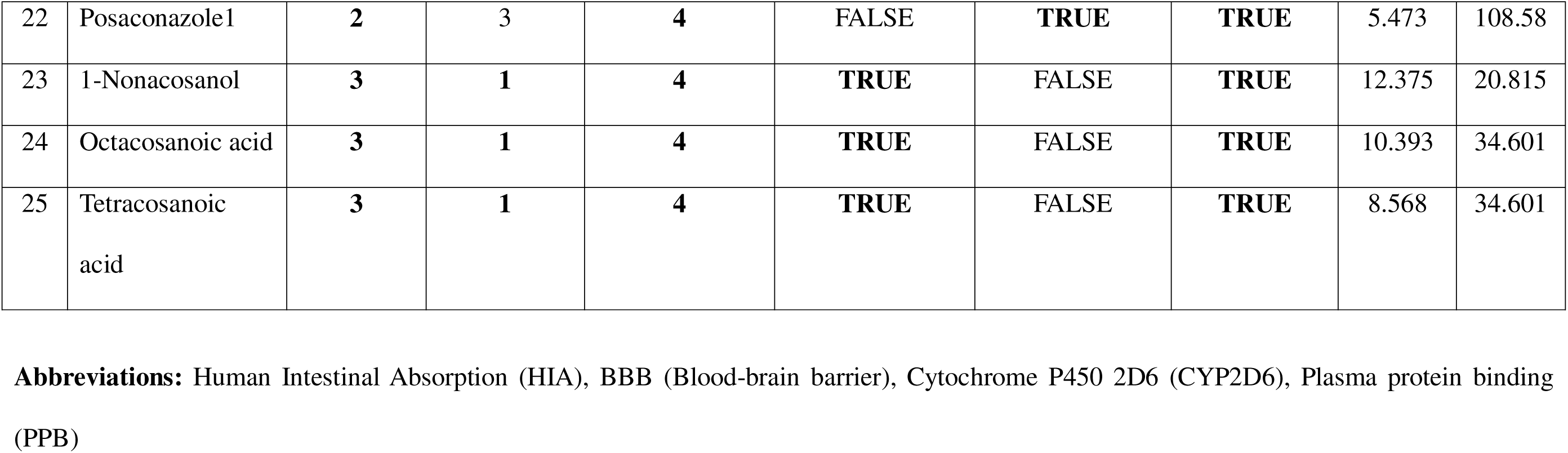
Result of ADMET parameters for the drugs which were able to bind.

### ADMET Parameters

Among the 25 ligands (6 drugs/drug derivatives + 19 phytochemicals), only 18 ligands showed satisfactory intestinal absorption and blood-brain barrier penetration, as evident from the ADMET plot (Fig. 6). The ADMET plot of the compounds was plotted between AlogP98 and 2D PSA of the compounds, and the ligands which fall within the ellipsoids include the first 18 (S. No. 1-18) ligands in the table presented under the figure. The exact parameters for all the 25 ligands are presented in table 8.

**Fig. 6.**
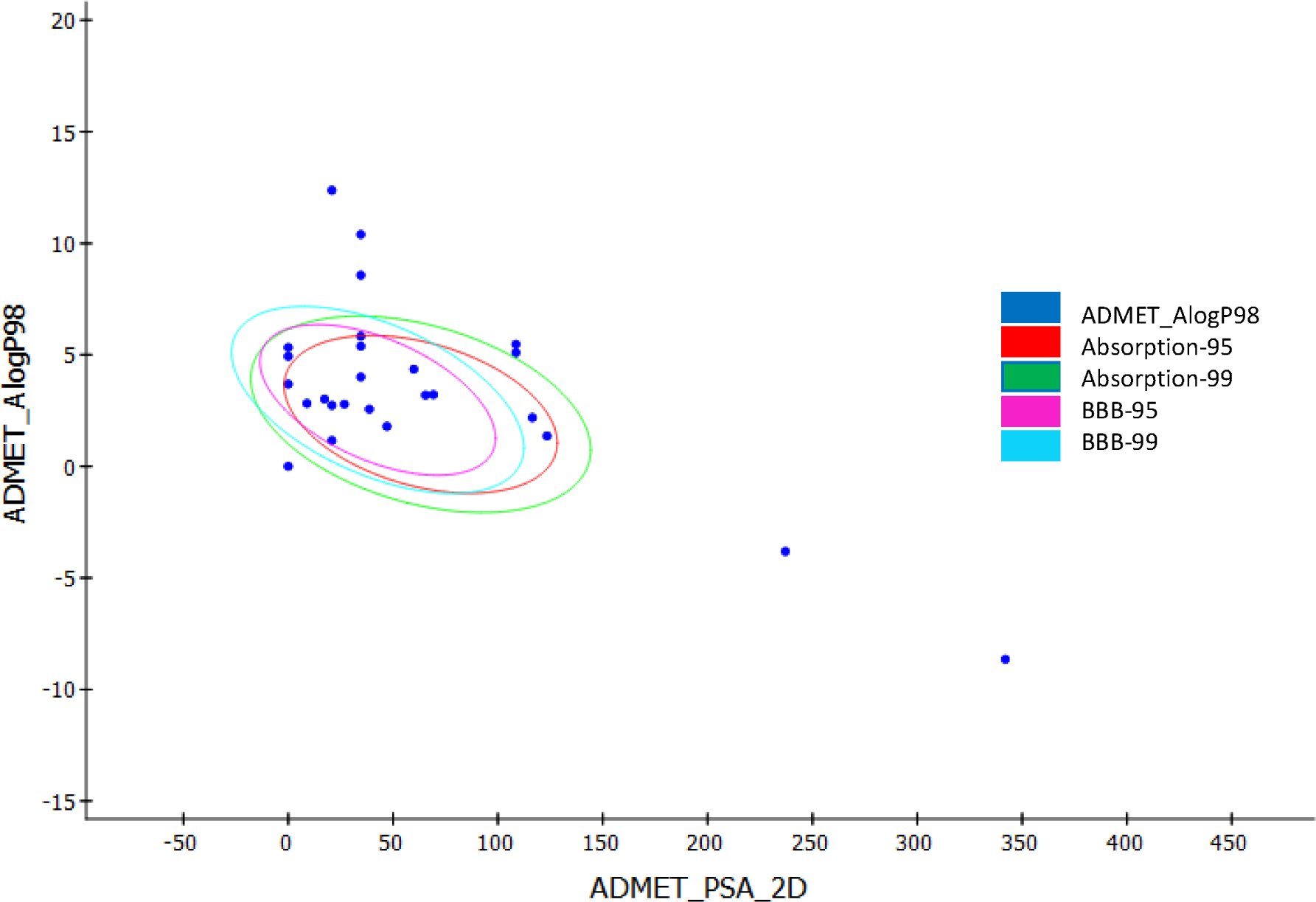
ADMET Plot

From table 8, it is evident that except 1-Nonacosanol, Octacosanoic acid and Tetracosanoic acid, all ligands show low to optimal aqueous solubility (Solubility level 2-4), and hence, greater drug likeliness. Except some ligands such as Posaconazole, Nikkomycin1, Polyoxin1, 1-Nonacosanol, Octacosanoic acid and Tetracosanoic acid, all other ligands show good to moderate human intestinal absorption (HIA level 0-1). 4-Allyl-2,6-dimethoxyphenol, β-Bisabolene, Citronellal, Decane, Dihydrocapsaicin, Elemicin, Estragole, Linalool, Myrcene, Myristic acid, Oleic acid, and Stearic acid possess good blood-brain barrier penetration ability (BBB level 0-1), and rest others have extremely low to undefined penetration. Some of the ligands such as Stearic acid, β-Bisabolene, 1-Nonacosanol, Octacosanoic acid and Tetracosanoic acid, etc. bind to plasma protein (PPB = True) which indicates its incapability to execute therapeutic action. Other parameters of the ligands such as hepatotoxicity and Cytochrome P450 2D6 (CYP2D6) inhibition can also be found from the table.

### TOPKAT Parameters

Table 9 shows that 4-Allyl-2,6-dimethoxyphenol, Cubebin, Dihydrocapsaicin, Zingerone, Nikkomycin1, and Polyoxin1 are the ligands that do not show any form of carcinogenicity in Male/Female Rat as well as Mouse models as per FDA parameters. Rest all ligands show some form of carcinogenicity as per the TOPKAT report. None of the ligands are reported to be mutagenic. Isavuconazole4, 1-Penten-3-ol, 4-Allyl-2, 6-dimethoxyphenol, β-Bisabolene, Elemicin, Estragole, Linalool, Myrcene, Zingerone, Posaconazole, Posaconazole1, and 1-Nonacosanol are reported to have developmental toxicity potential. Decane was shown to possess severe skin irritancy, whereas 1-Penten-3-ol, 4-Allyl-2, 6-dimethoxyphenol, Citronellal, Cubebin, Linalool and Nikkomycin1 are reported to possess severe ocular irritancy. Out of all these ligands, Dihydrocapsaicin and Polyoxin1 are shown to be non-carcinogenic, non-mutagenic, non-toxic, and possess mild to moderate skin and ocular irritancy. Apart from this, Cubebin and Nikkomycin1 prove to be promising ligands, with severe ocular irritancy as a disadvantageous effect.

**Table 9.**
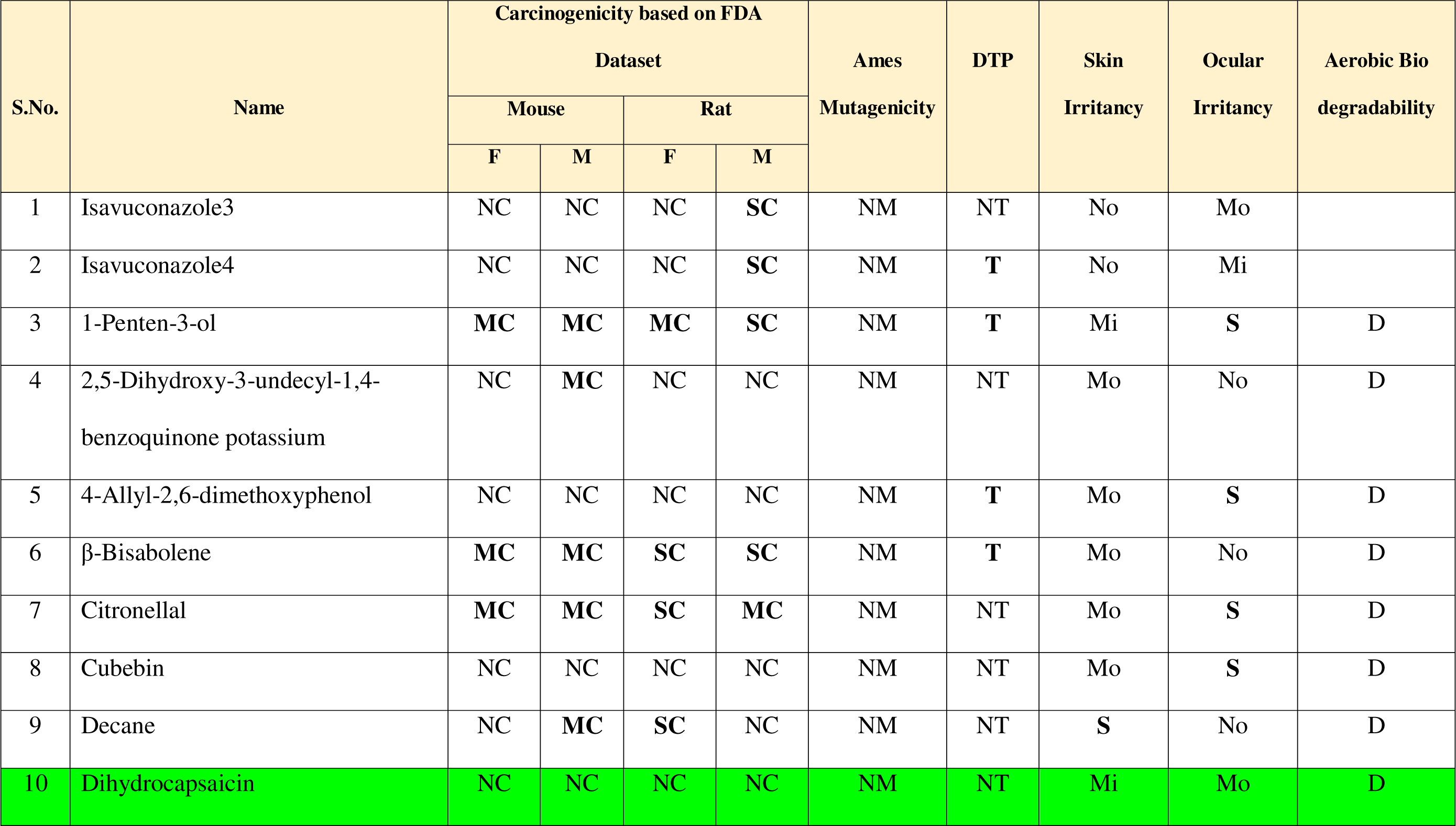
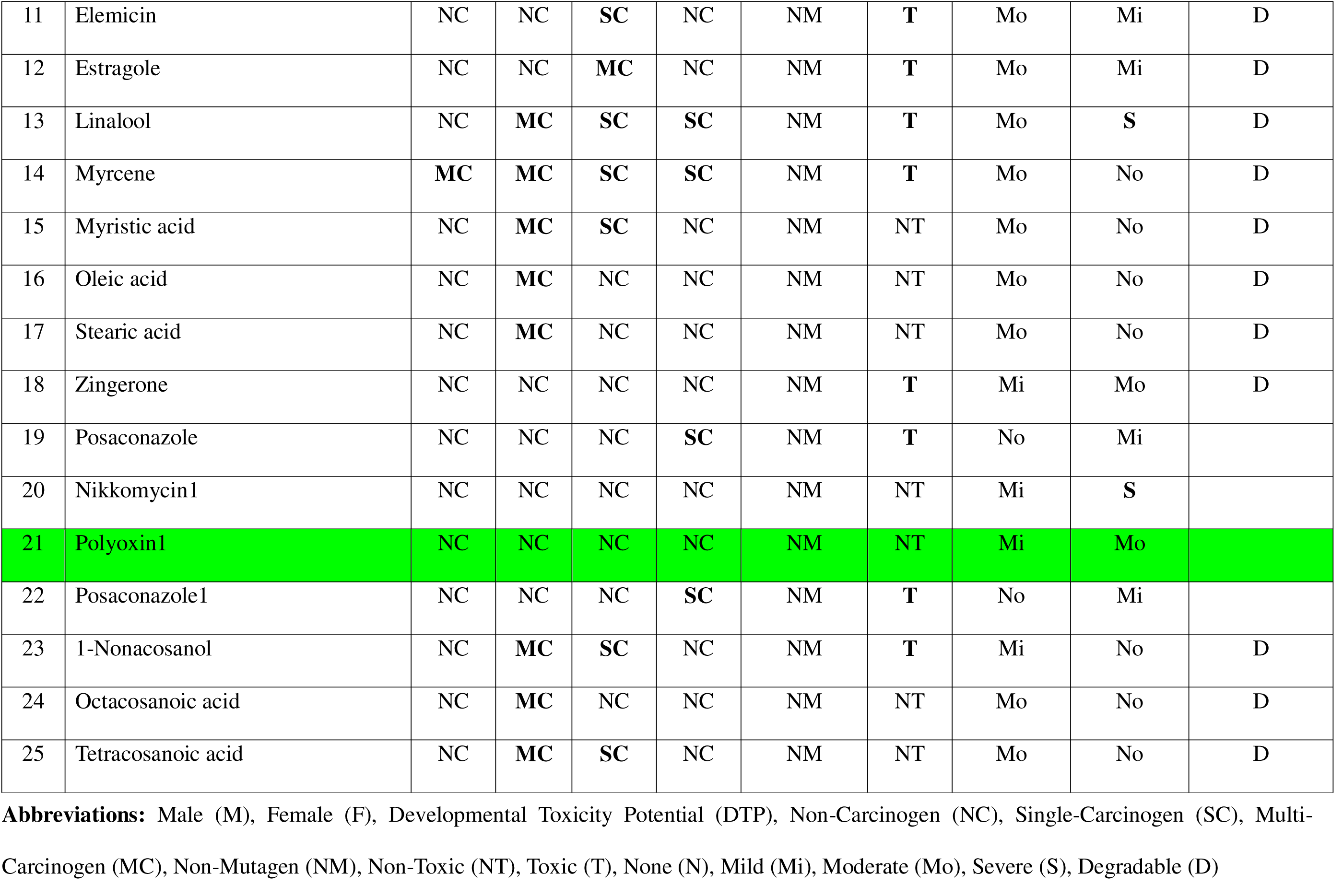
TOPKAT parameters for the ligands which were able to bind the protein.

### Compliance for Lipinski and Veber’s Rules

Table 10 shows molecular properties of ligands which pass Lipinski’s and Veber’s rule. Among the 25 ligands which were able to bind to the protein, only 15 complied with the Lipinski rule while 10 drugs failed. Modified drug derivatives Isavuconazole3 and Isavuconazole4 are found to comply with the Lipinski rule. Rest all molecules which are found to pass the screening are the plant derived anti-fungal ligands. It is surprising to note that top binding modified drug derivatives are found to fail the Lipinski rule.

**Table 10.** Different molecular parameters for the ligands which pass the Lipinski and Veber’s rules.

| S. No. | Name | Mol. Wt. | ALogP | RB | PSA | HBA | HBD |
| --- | --- | --- | --- | --- | --- | --- | --- |
| 1 | Isavuconazole3 | 462.493 | 2.61 | 6 | 146.57 | 8 | 3 |
| 2 | Isavuconazole4 | 496.508 | 1.309 | 7 | 153.76 | 10 | 4 |
| 3 | 1-Penten-3-ol | 86.1323 | 1.159 | 2 | 20.23 | 1 | 1 |
| 4 | 2,5-Dihydroxy-3-undecyl-1,4-benzoquinone potassium | 292.37 | 3.214 | 10 | 80.25 | 4 | 0 |
| 5 | 4-Allyl-2,6-dimethoxyphenol | 194.227 | 2.563 | 4 | 38.69 | 3 | 1 |
| 6 | β-Bisabolene | 204.351 | 5.329 | 4 | 0 | 0 | 0 |
| 7 | Citronellal | 154.249 | 3.018 | 5 | 17.07 | 1 | 0 |
| 8 | Cubebin | 356.369 | 3.194 | 4 | 66.38 | 6 | 1 |
| 9 | Decane | 142.282 | 4.933 | 7 | 0 | 0 | 0 |
| 10 | Dihydrocapsaicin | 307.428 | 4.354 | 10 | 58.56 | 4 | 2 |
| 11 | Elemicin | 208.254 | 2.788 | 5 | 27.69 | 3 | 0 |
| 12 | Estragole | 148.202 | 2.821 | 3 | 9.23 | 1 | 0 |
| 13 | Linalool | 154.249 | 2.735 | 4 | 20.23 | 1 | 1 |
| 14 | Myrcene | 136.234 | 3.687 | 4 | 0 | 0 | 0 |
| 15 | Zingerone | 194.227 | 1.792 | 4 | 46.53 | 3 | 1 |

**Abbreviations:** Rotatable bonds (RB), Molecular Polar Surface Area (PSA), Hydrogen bond acceptors (HBA), Hydrogen bond donors (HBD)

### Interaction studies of the top 15 ligands with the protein

The docked complex of each of the top 15 ligands (sorted after ADMET analysis, and Lipinski rule compliance) in the receptor cavity and the interacting residues are shown in Fig. 7(a) to (o)

**Fig. 7(a).**
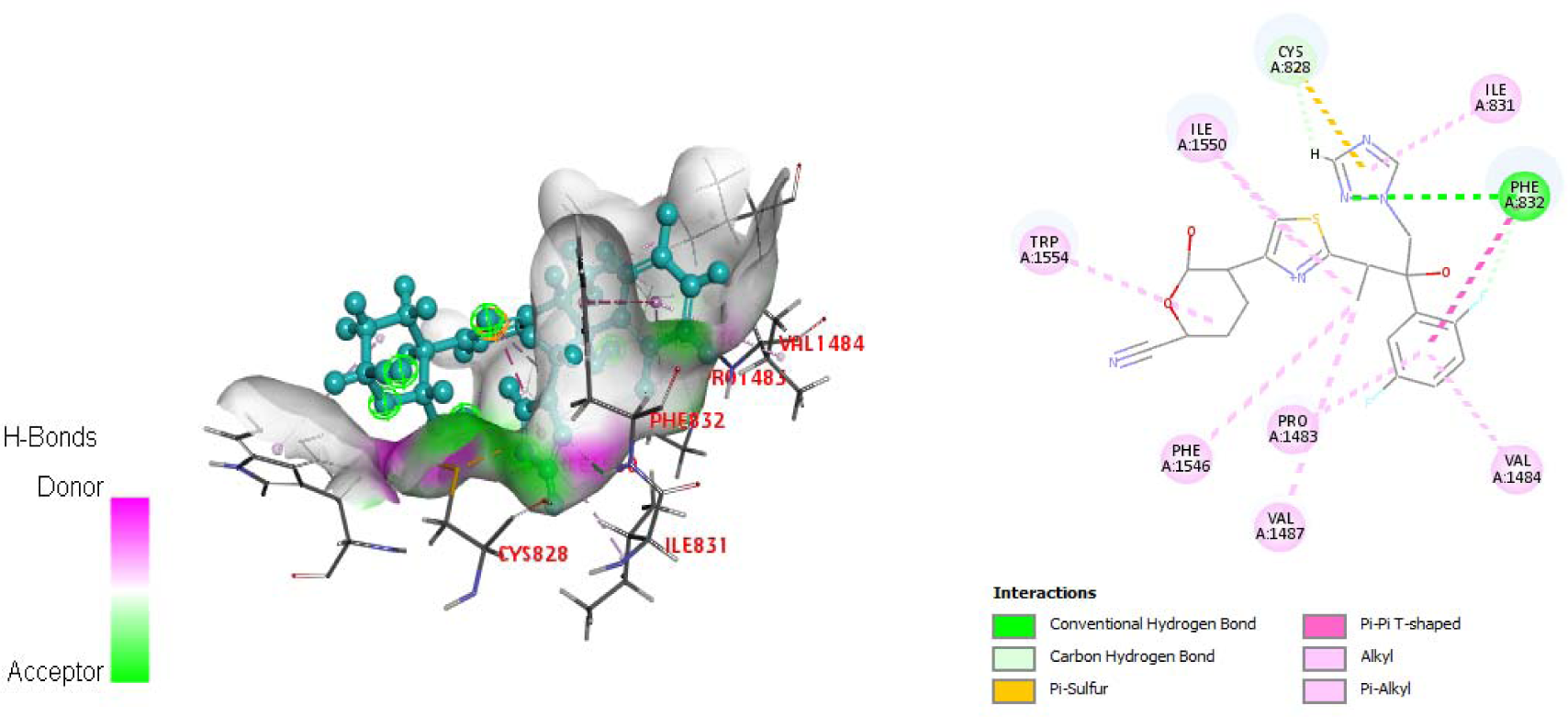
Modelled Chitin Synthase docked with Isavuconazole3; Key interacting residues: Cys828, Ile831, Phe832, Pro1483, Val1484, Val1487, Phe1546, Ile1550, Trp1554

**Fig. 7(b).**
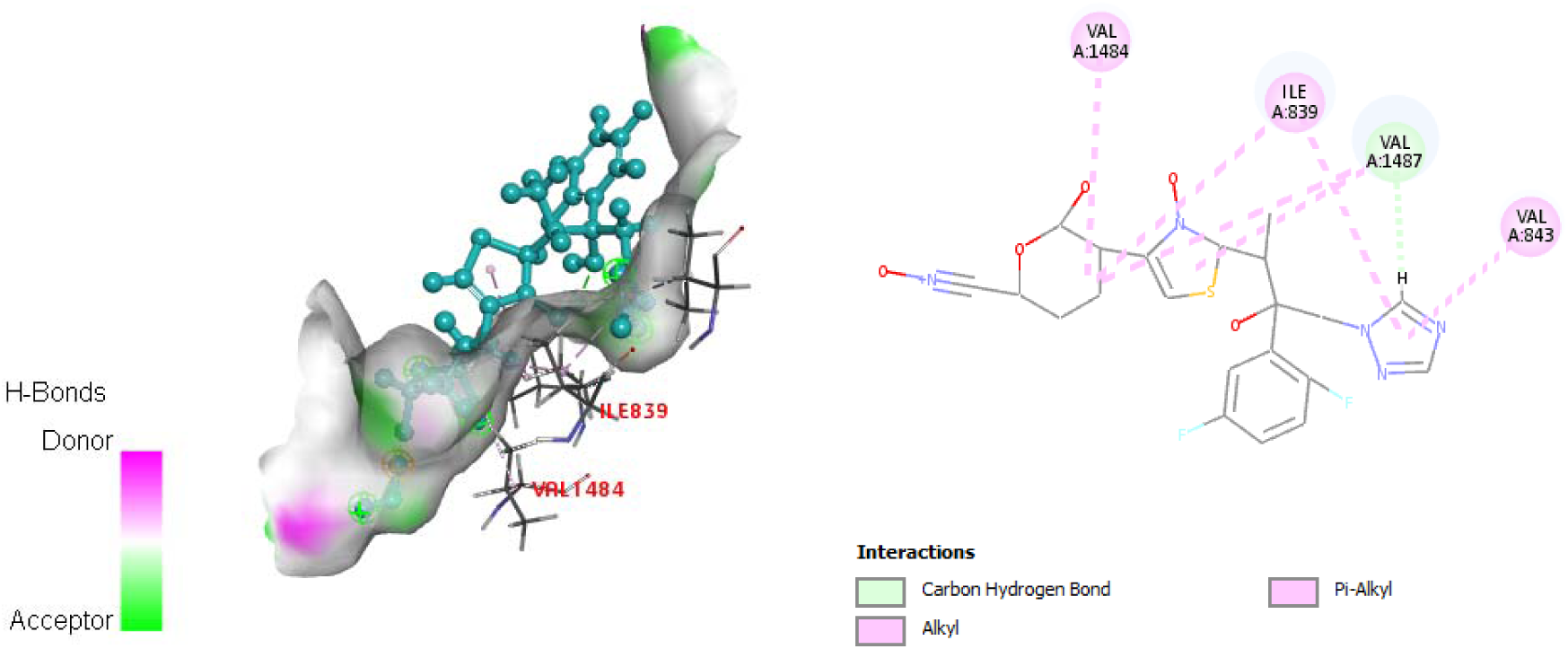
Modelled Chitin Synthase docked with Isavuconazole4; Key interacting residues: Ile839, Val843, Val1484, Val1487

**Fig. 7(c).**
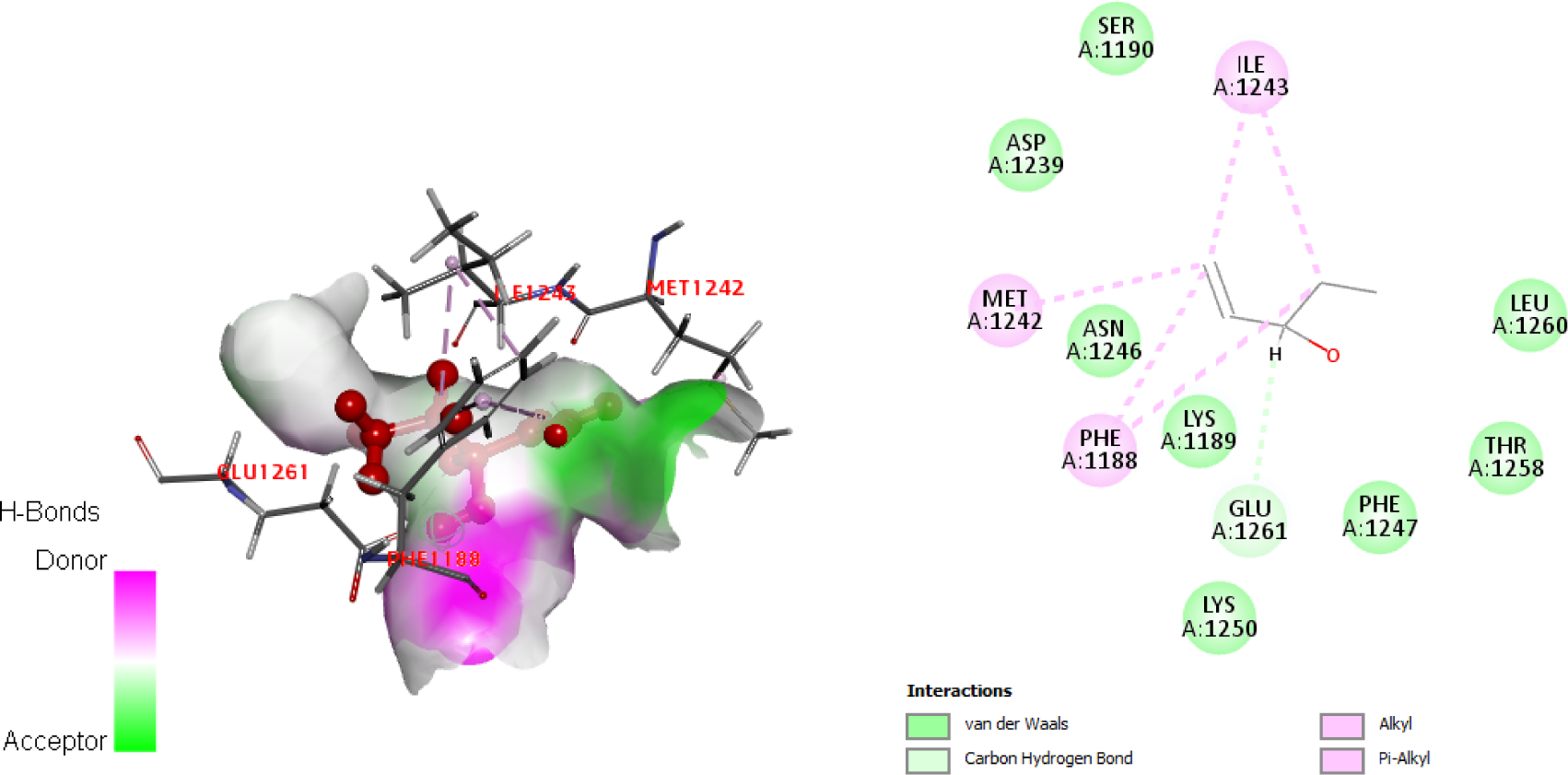
Modelled Chitin Synthase docked with 1-penten-3-ol; Key interacting residues: Phe1188, Lys1189, Ser1190, Asp1239, Met1242, Asn1246, Phe1247, Lys1250, Thr1258, Leu1260, Glu1261

**Fig. 7(d).**
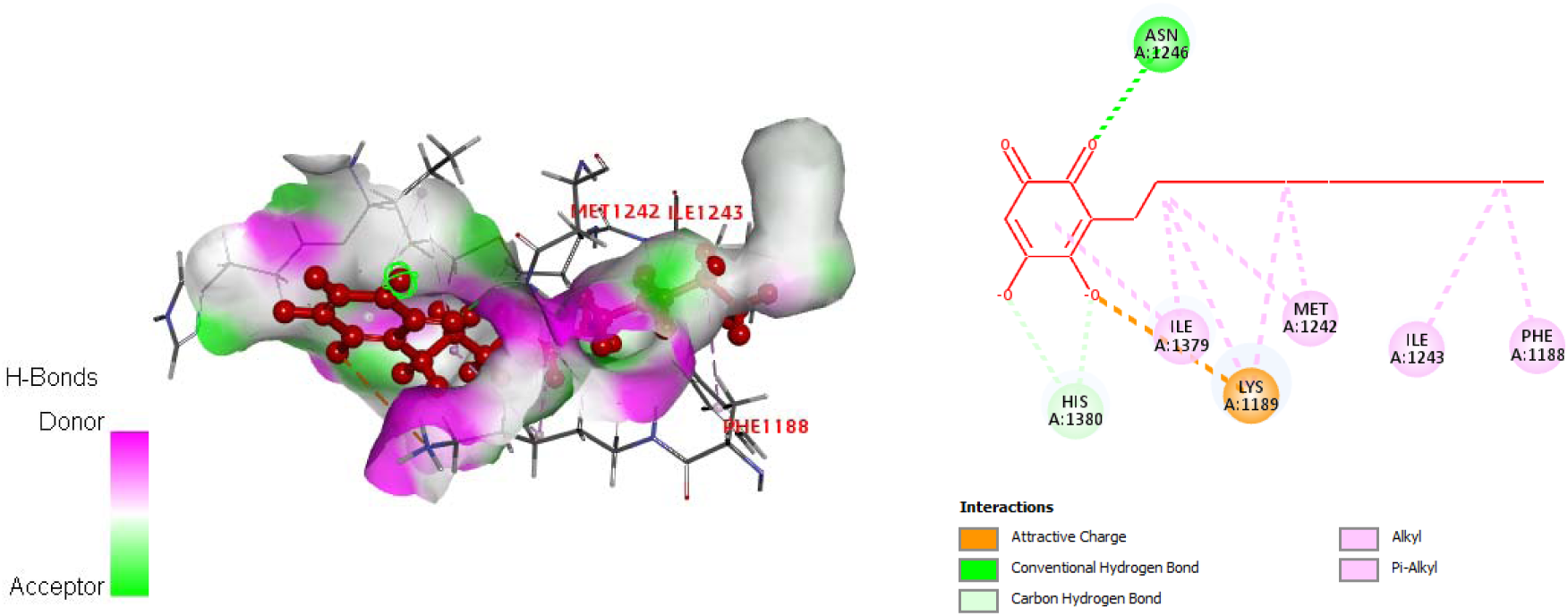
Modelled Chitin Synthase docked with 2,5-Dihydroxy-3-undecyl-1,4-benzoquinone potassium; Key interacting residues: Phe1188, Lys1189, Met1242, Ile1243, Asn1246, Ile1379, His1380

**Fig. 7(e).**
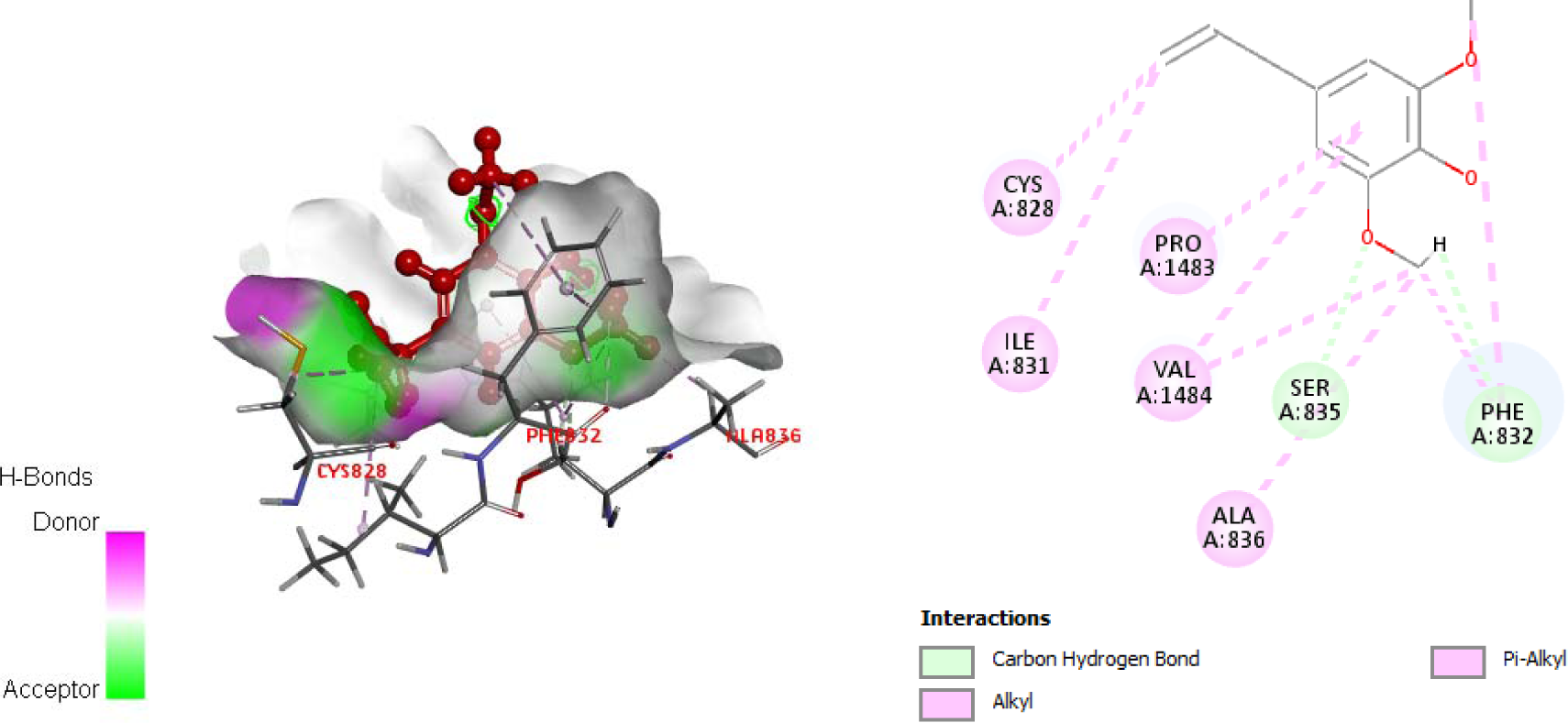
Modelled Chitin Synthase docked with 4-Allyl-2,6-dimethoxyphenol; Key interacting residues: Cys828, Ile831, Phe832, Ser835, Ala836, Pro1483, Val1484

**Fig. 7(f).**
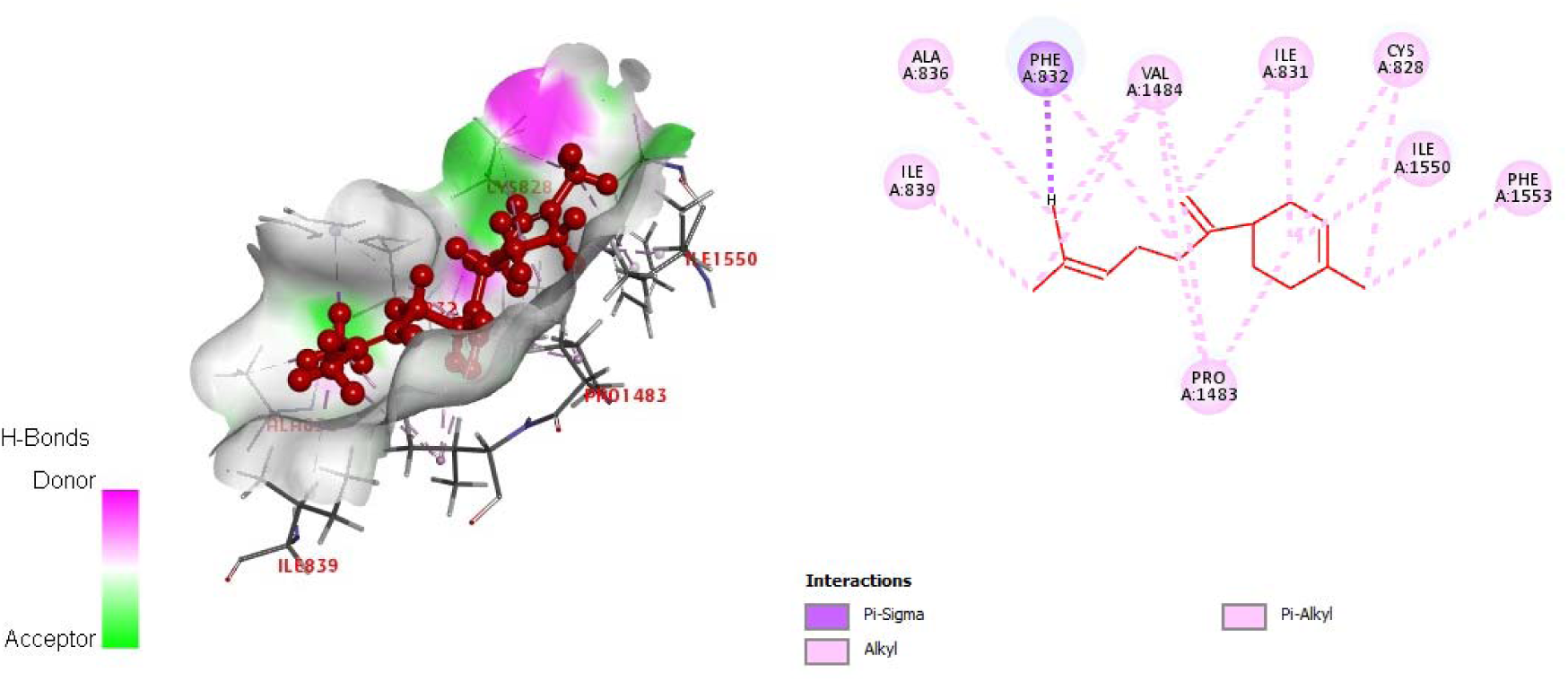
Modelled Chitin Synthase docked with β-Bisabolene; Key interacting residues: Cys828, Ile831, Phe832, Ser835, Ala836, Ile839, Pro1483, Val1484, Ile1550, Phe1553

**Fig. 7(g).**
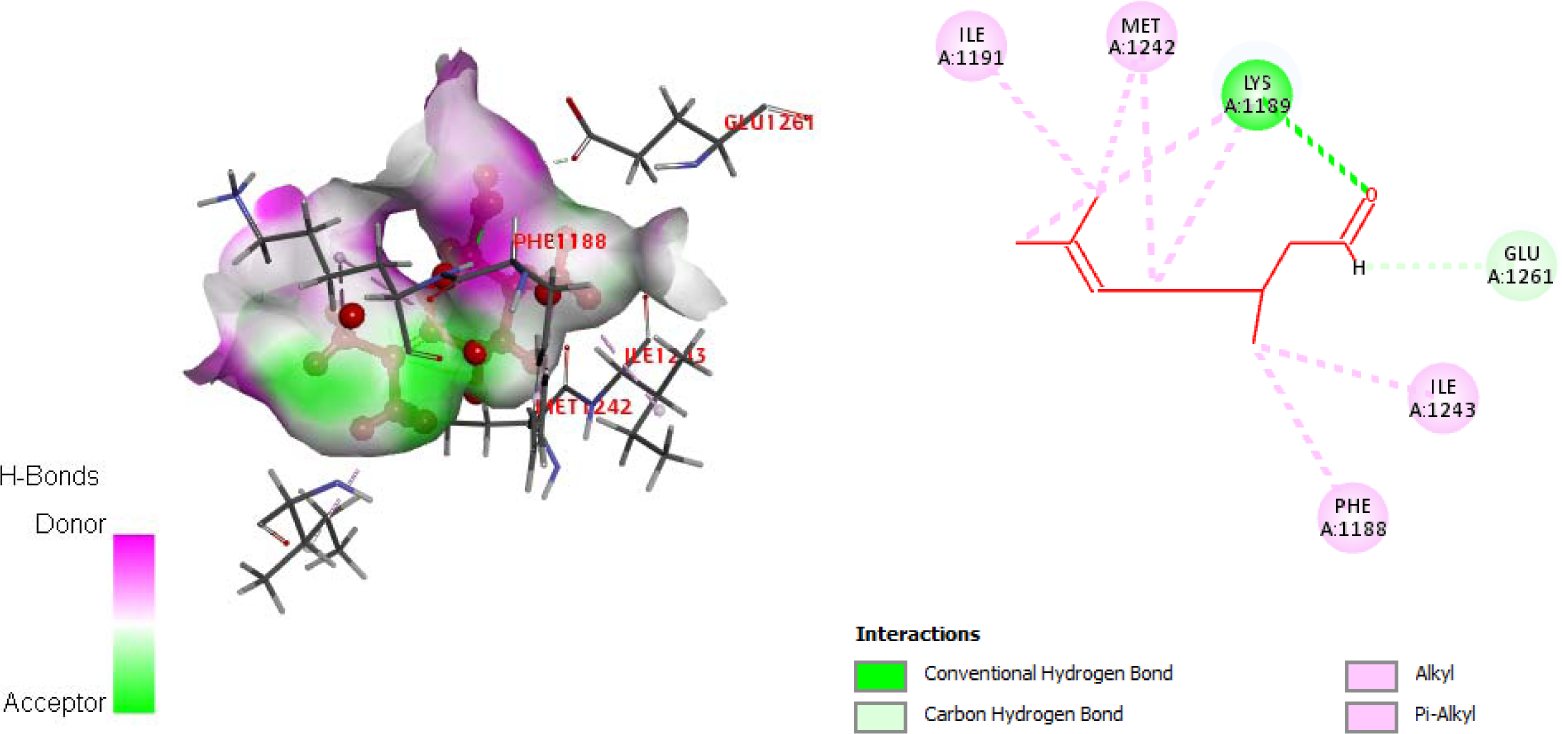
Modelled Chitin Synthase docked with Citronellal; Key interacting residues: Phe1188, Lys1189, Ile1191, Met1242, Ile1243, Glu1261

**Fig. 7(h).**
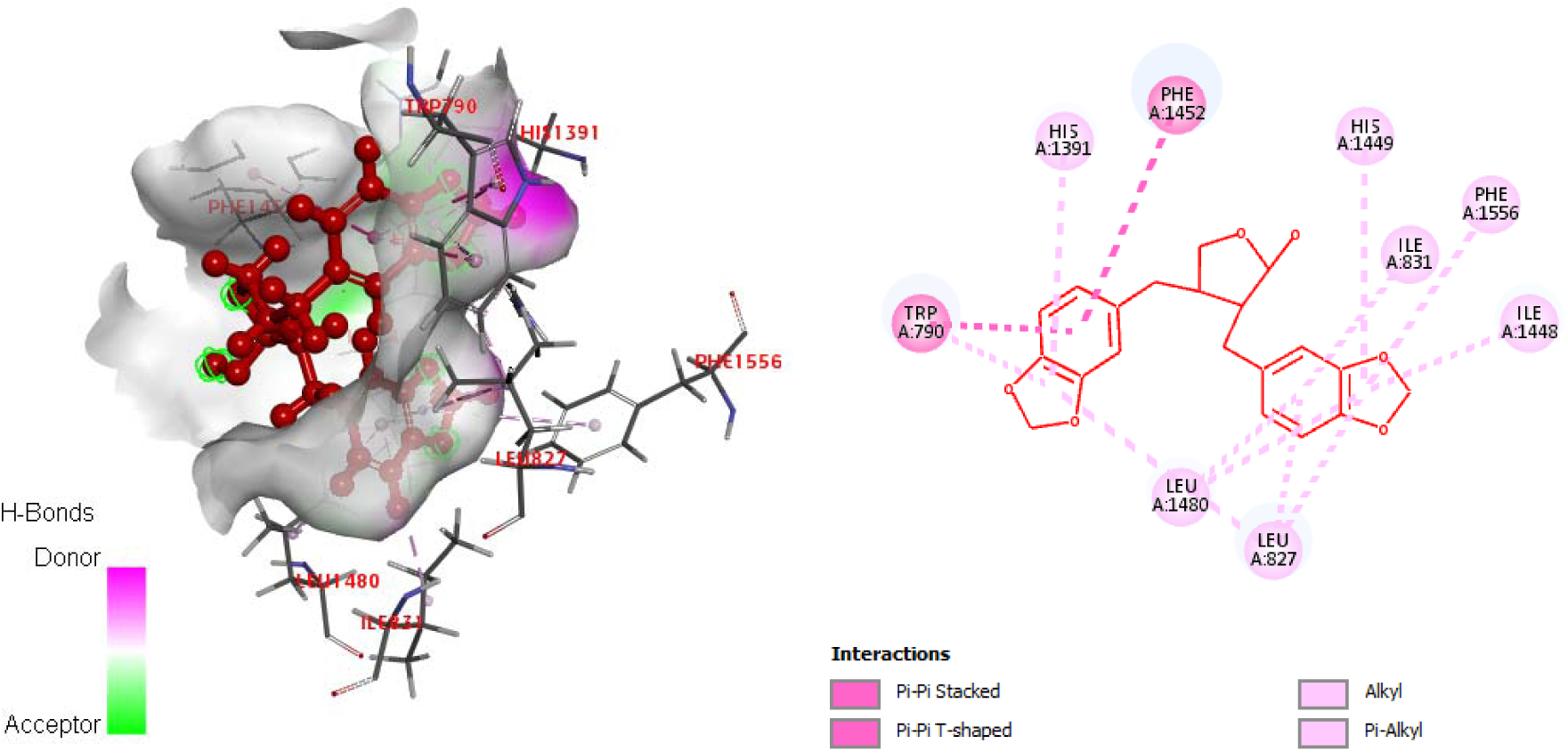
Modelled Chitin Synthase docked with Cubebin; Key interacting residues: Trp790, Leu827, Ile831, His1391, Ile1448, His1449, Phe1452, Leu1480, Phe1556

**Fig. 7(i).**
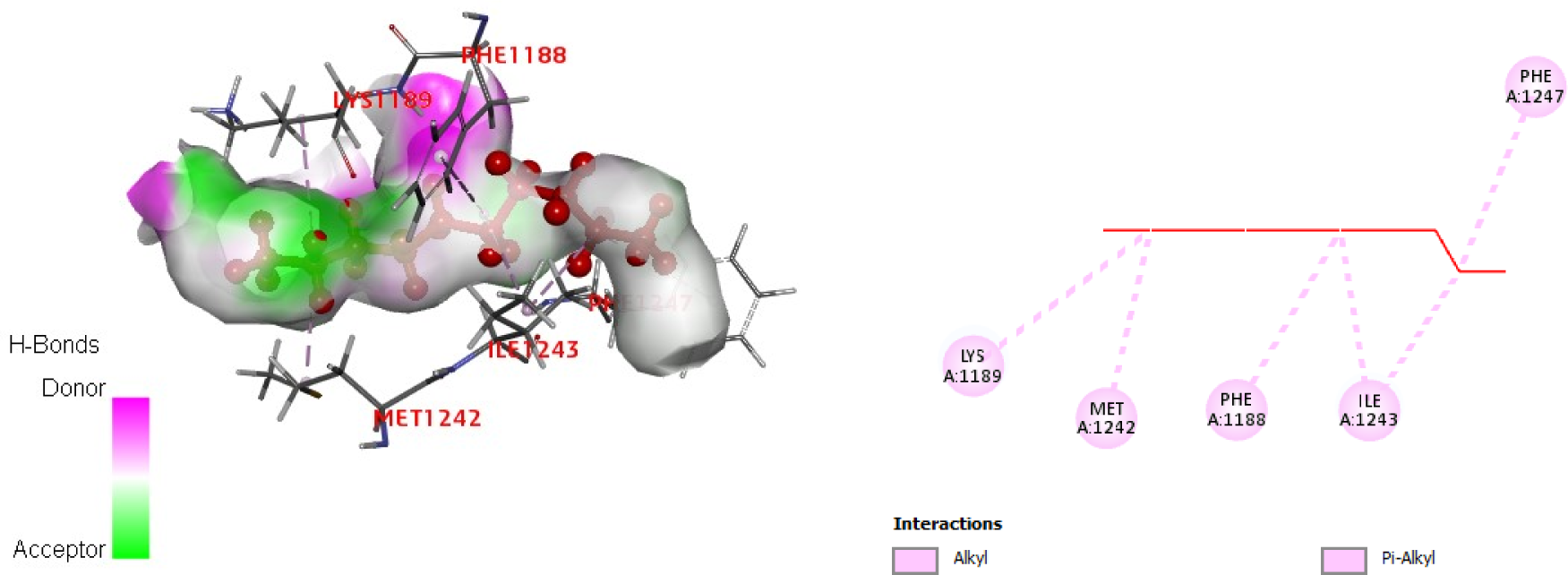
Modelled Chitin Synthase docked with Decane; Key interacting residues: Phe1188, Lys1189, Met1242, Ile1243, Phe1247

**Fig. 7(j).**
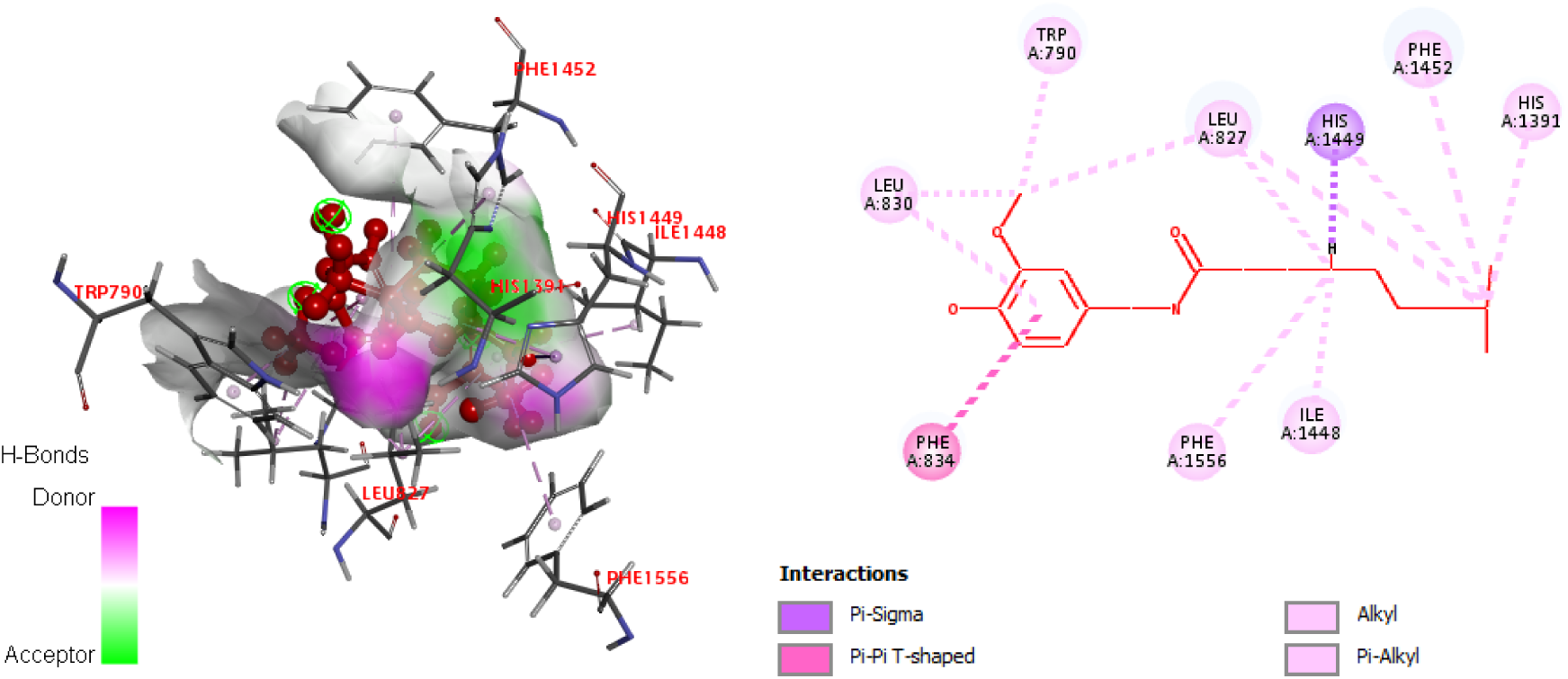
Modelled Chitin Synthase docked with Dihydrocapsaicin; Key interacting residues: Trp790, Leu827, Leu830, Phe834, His1391, Ile1448, His1449, Phe1452, Leu1480, Phe1556

**Fig. 7(k).**
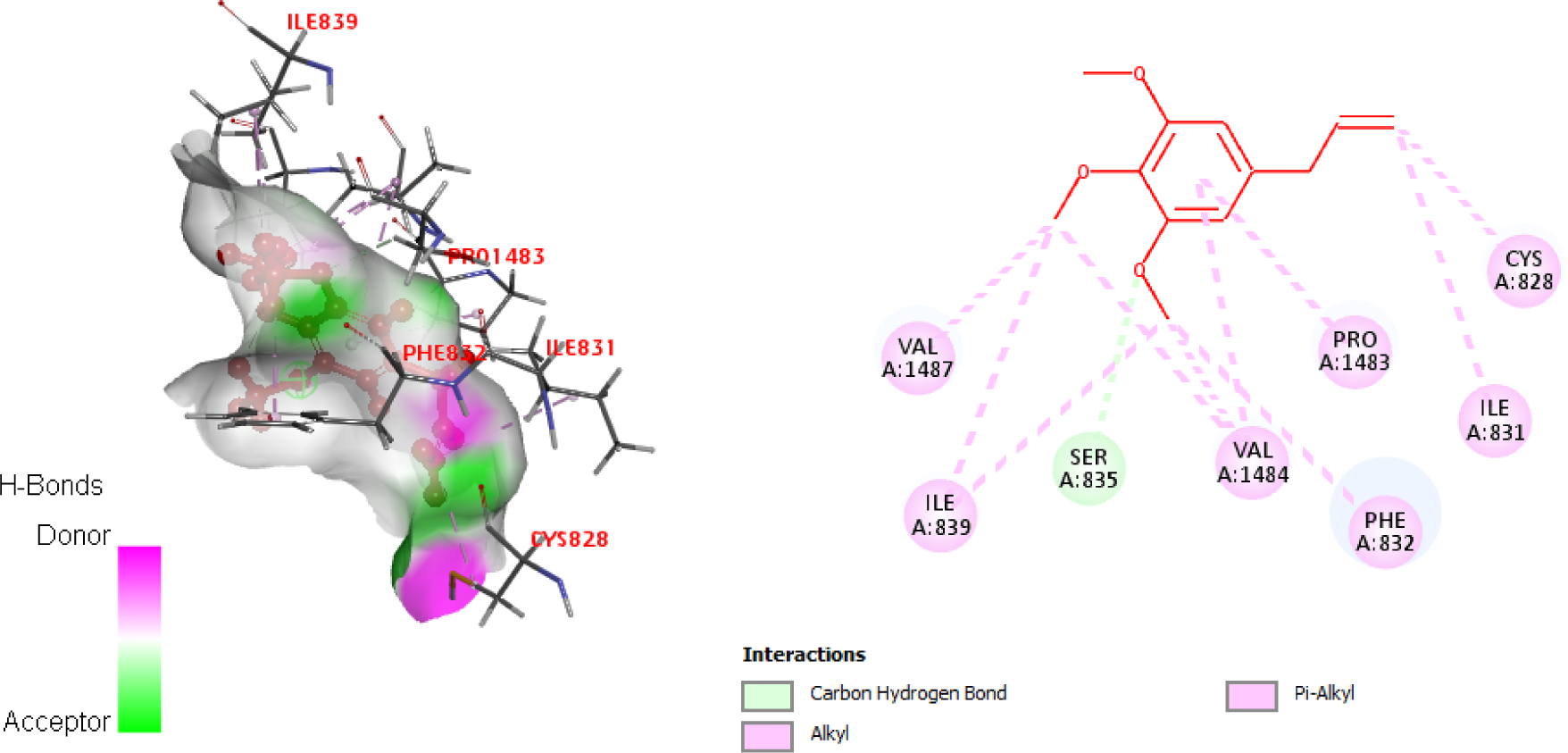
Modelled Chitin Synthase docked with Elemicin; Key interacting residues: Cys828, Ile831, Phe832, Ser835, Ile839, Pro1483, Val1484, Val1487

**Fig. 7(l).**
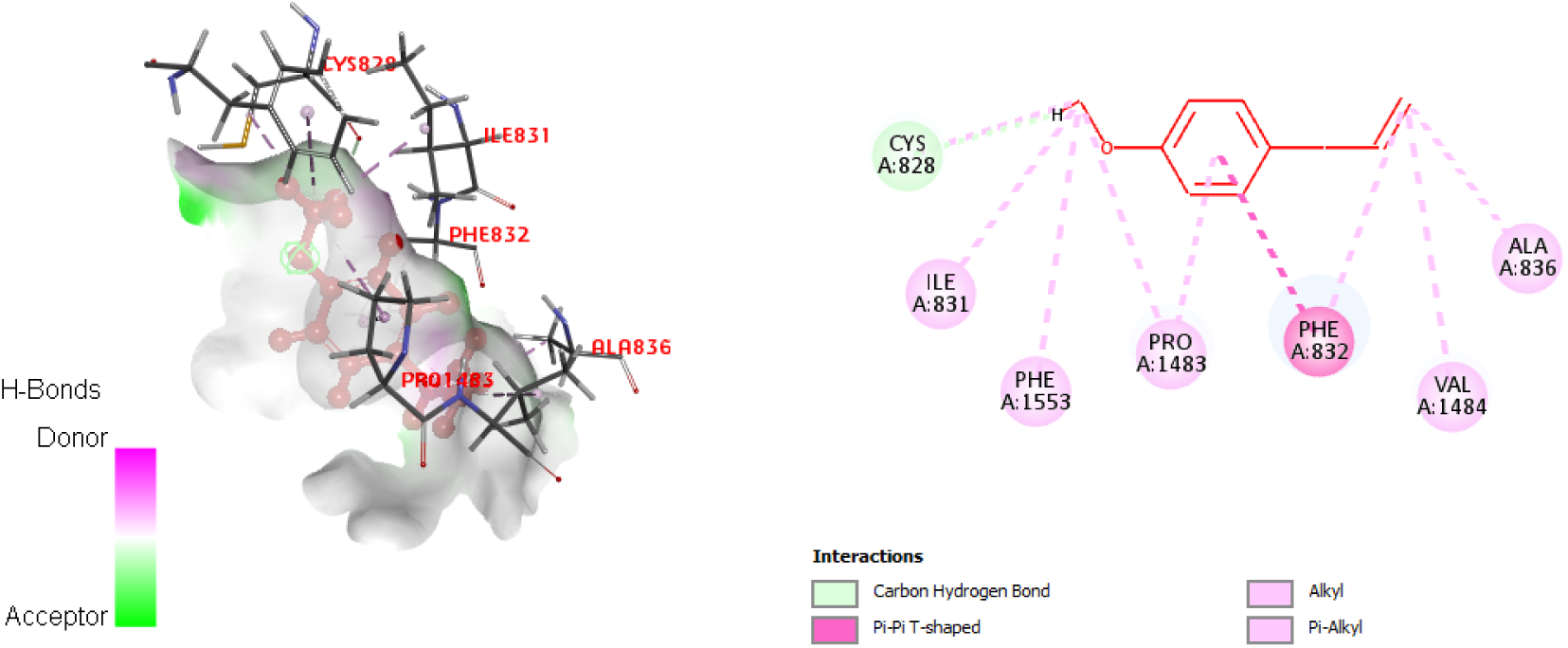
Modelled Chitin Synthase docked with Estragole; Key interacting residues: Cys828, Ile831, Phe832, Ala836, Ile839, Pro1483, Val1484, Phe1553

**Fig. 7(m).**
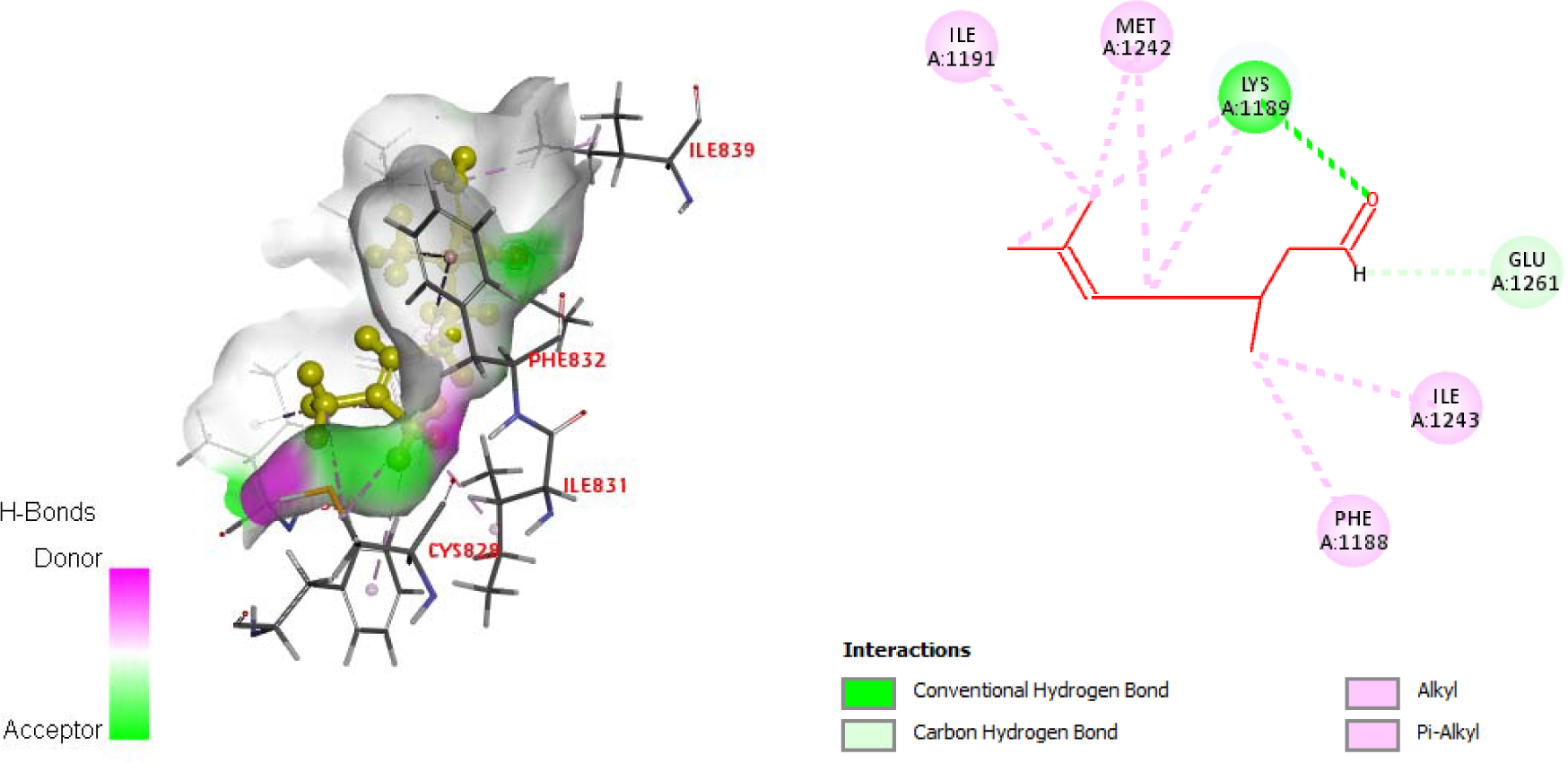
Modelled Chitin Synthase docked with Linalool; Key interacting residues: Phe1188, Lys1189, Ile1191, Met1242, Ile1243, Glu1261

**Fig. 7(n).**
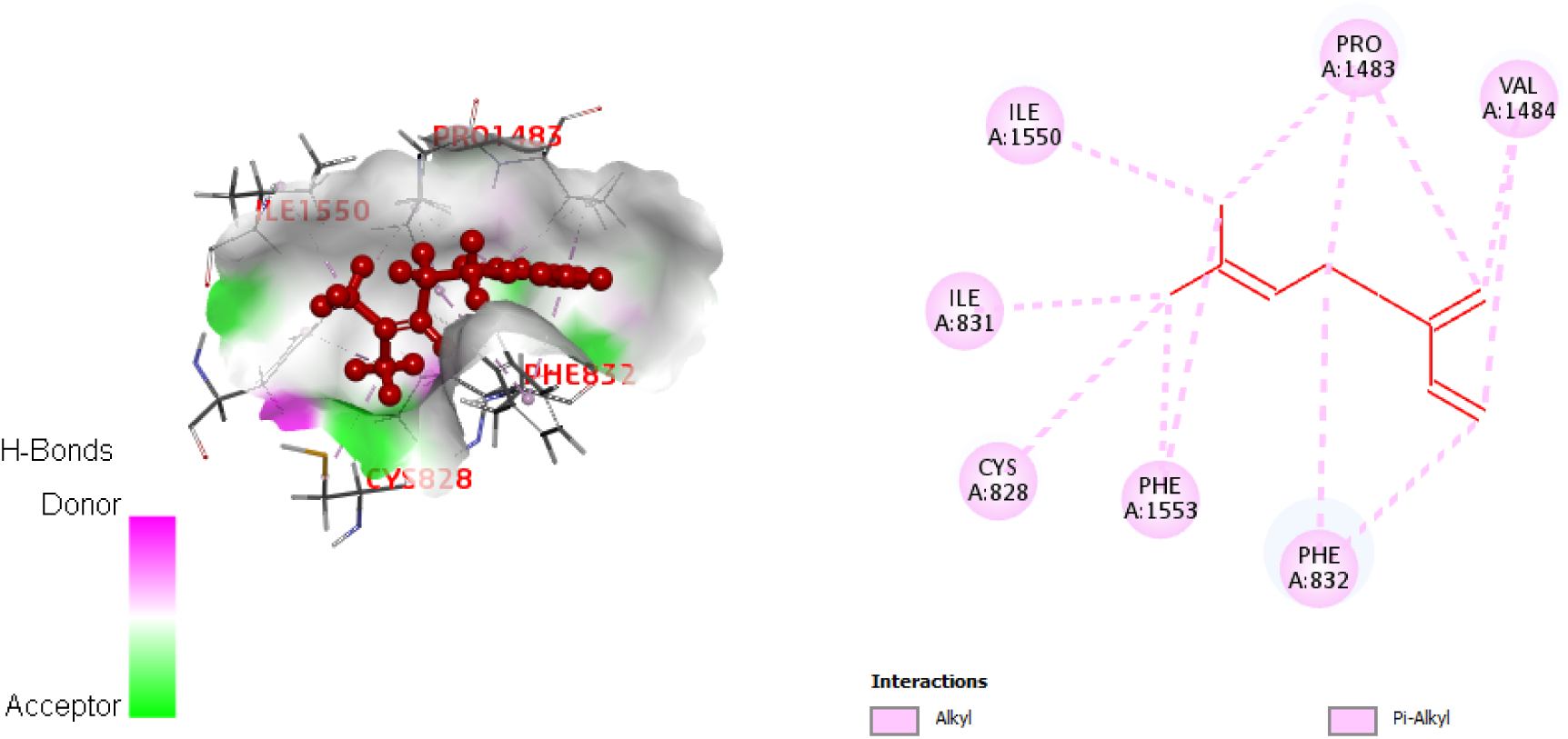
Modelled Chitin Synthase docked with Myrcene; Key interacting residues: Cys828, Ile831, Phe832, Pro1483, Val1484, Ile1550, Phe1553

**Fig. 7(o).**
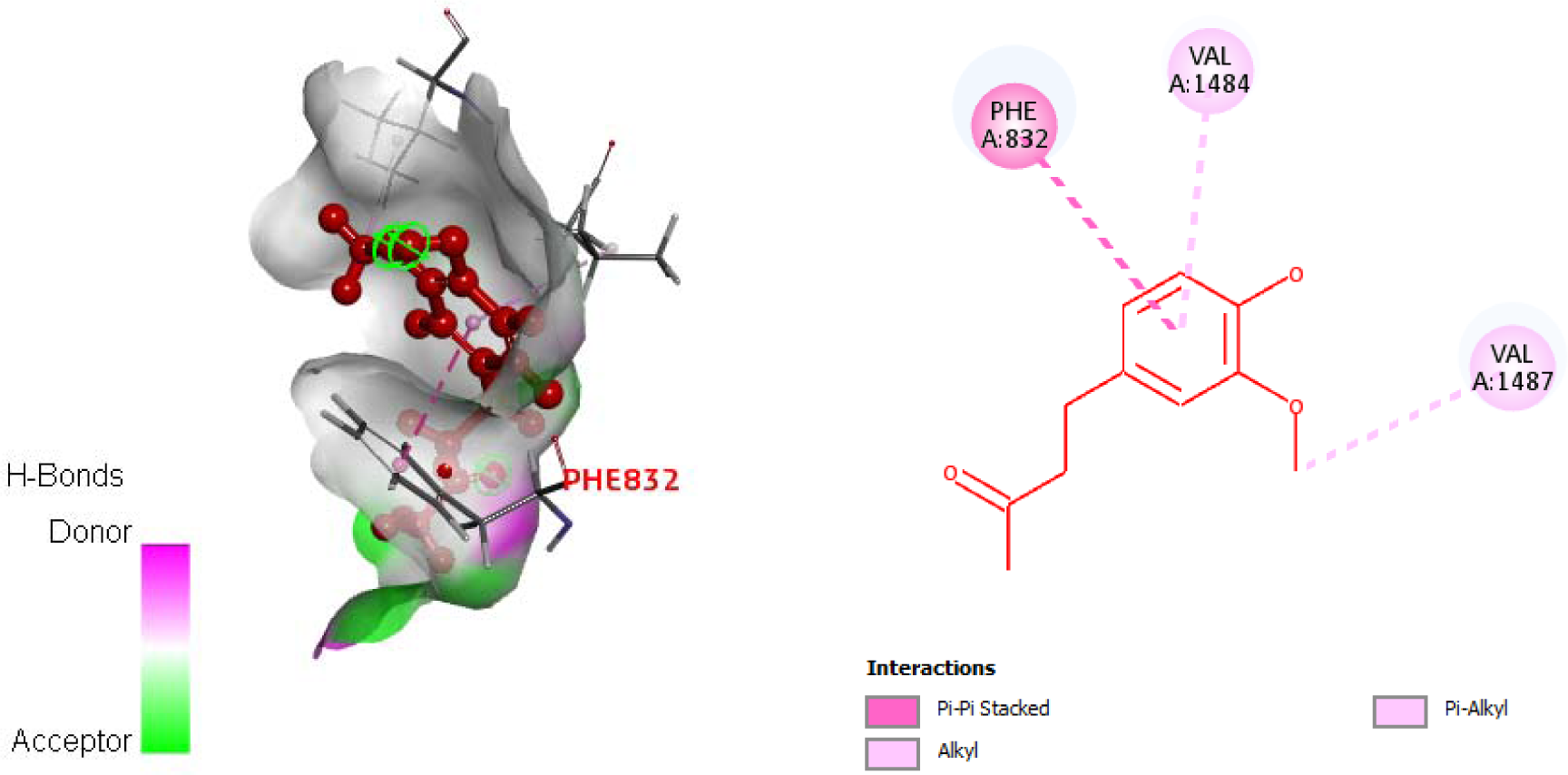
Modelled Chitin Synthase docked with Zingerone; Key interacting residues: Phe832, Val1484, Val1487

The key residues in the protein interacting with the ligands were identified as Ile 831, Cys828, Phe832, Pro1483, Val1484.

Table 11 shows the molecular interactions between the active site residues and the ligands.

**Table 11.**
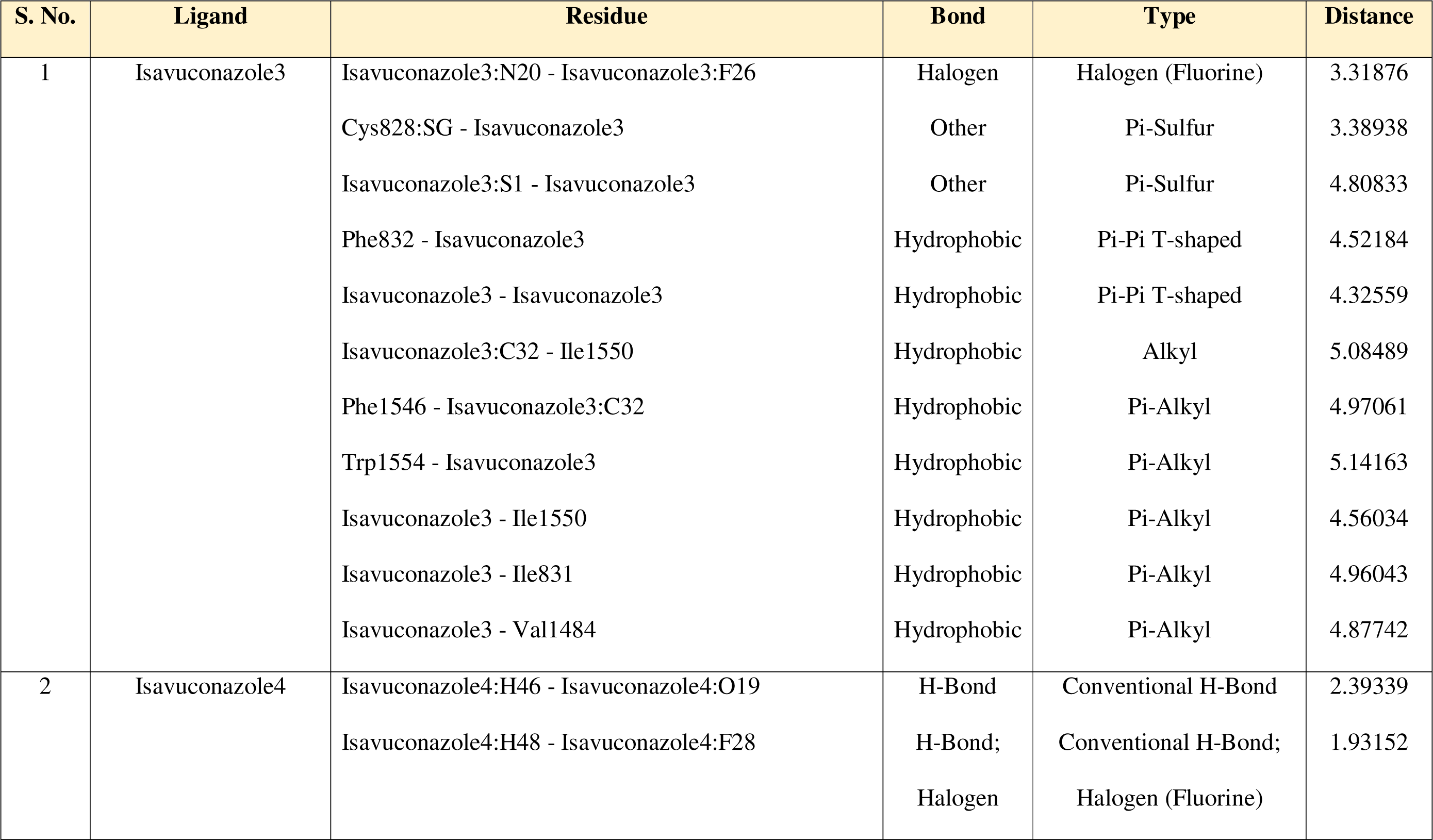
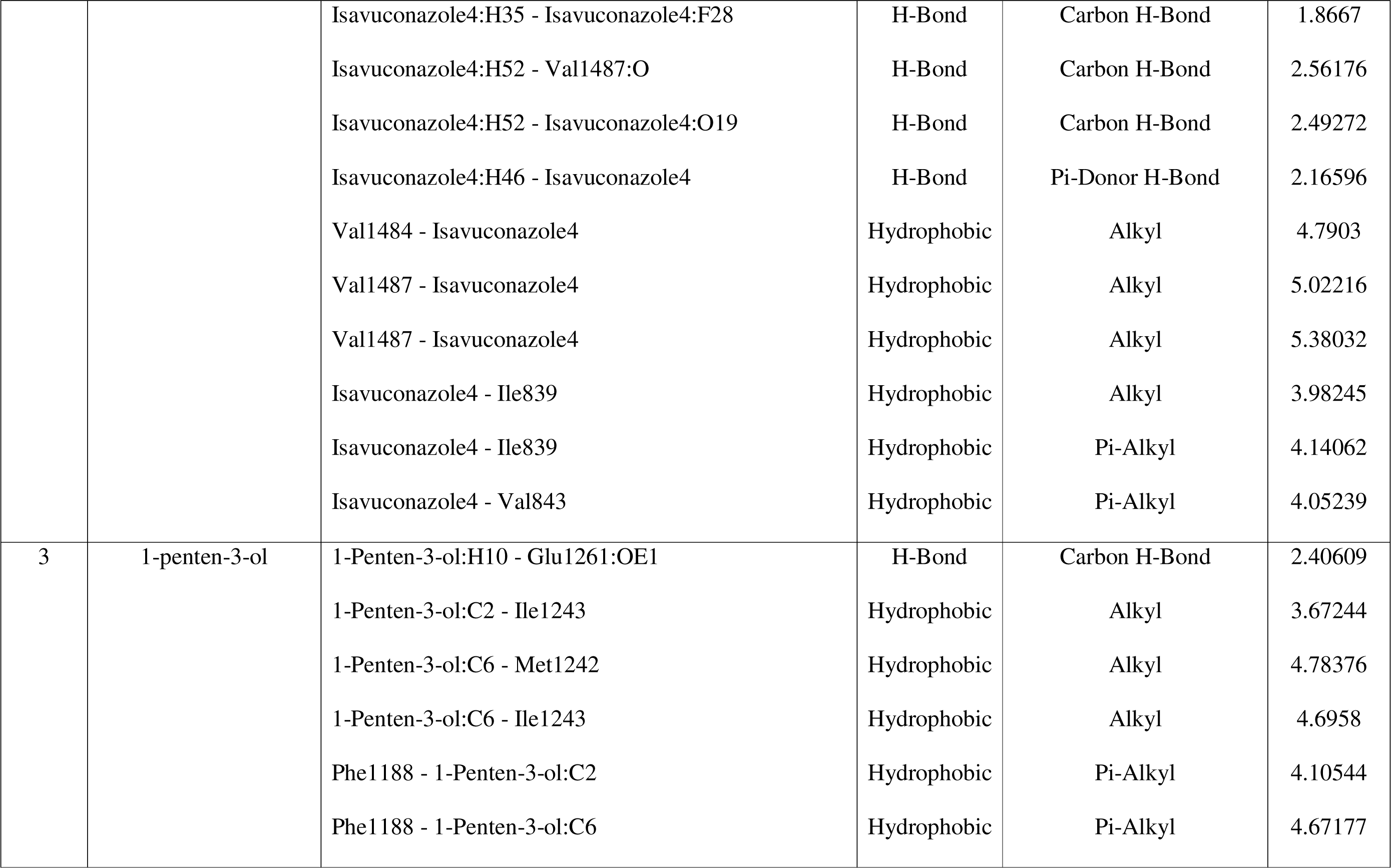
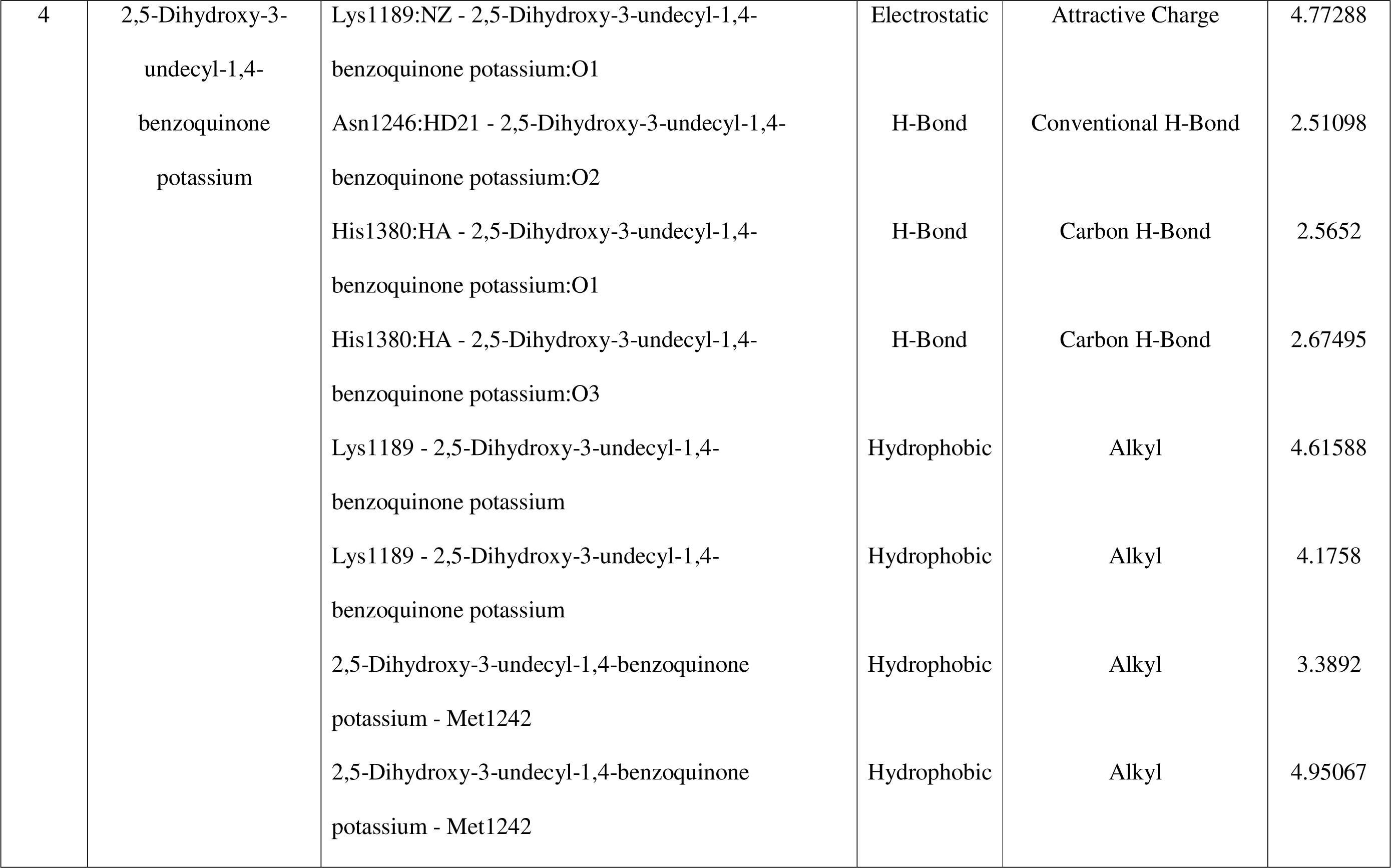
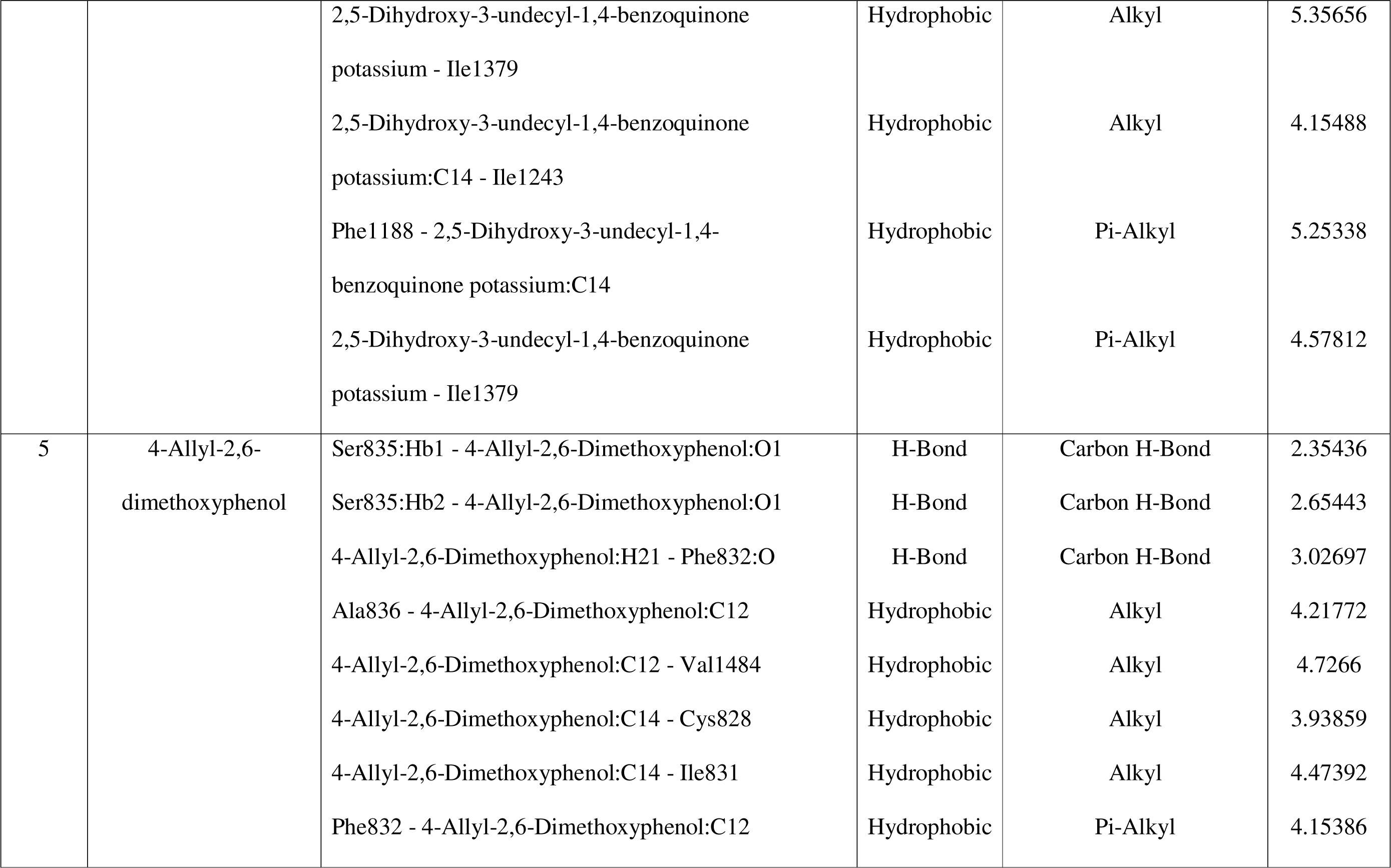
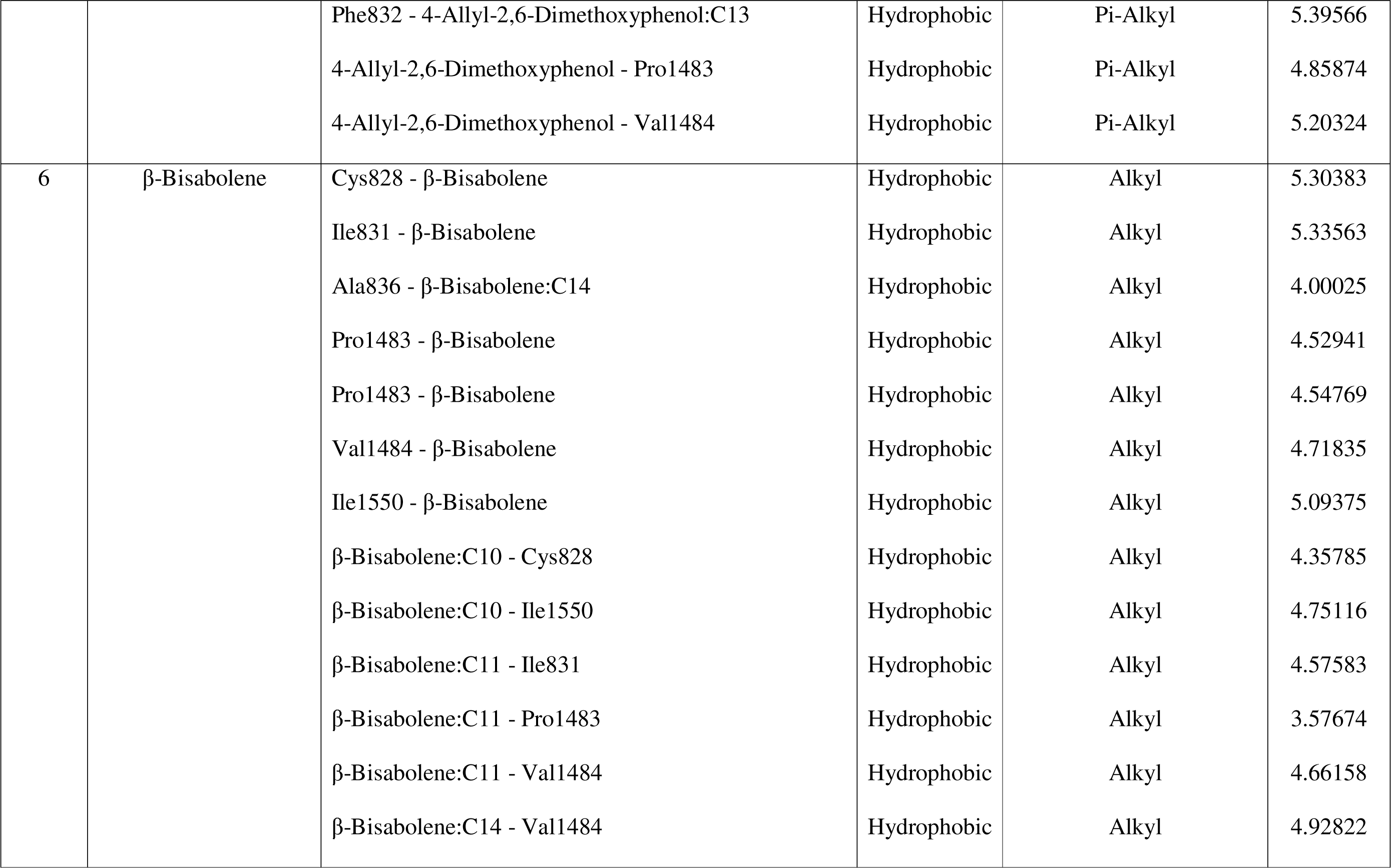
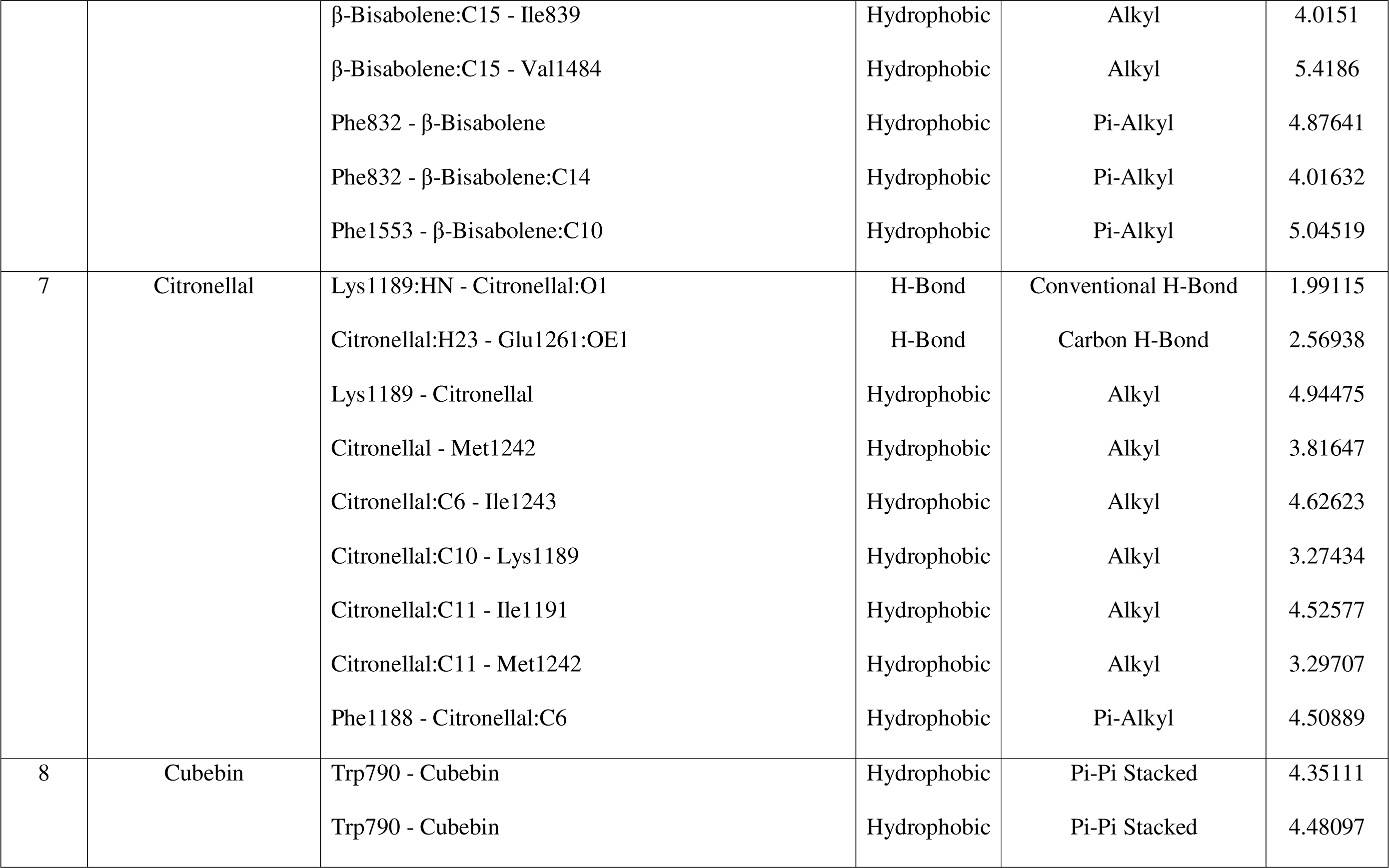
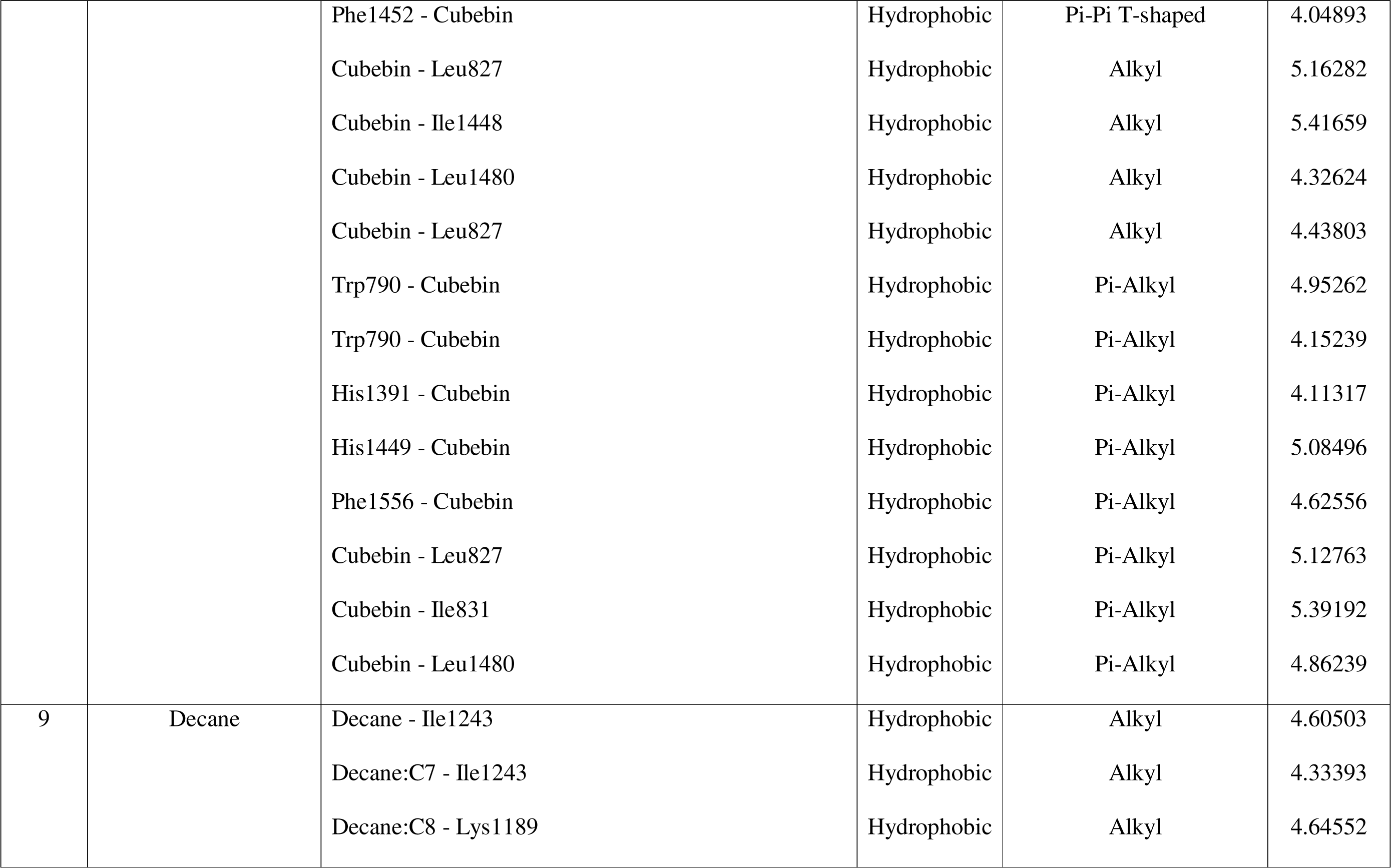
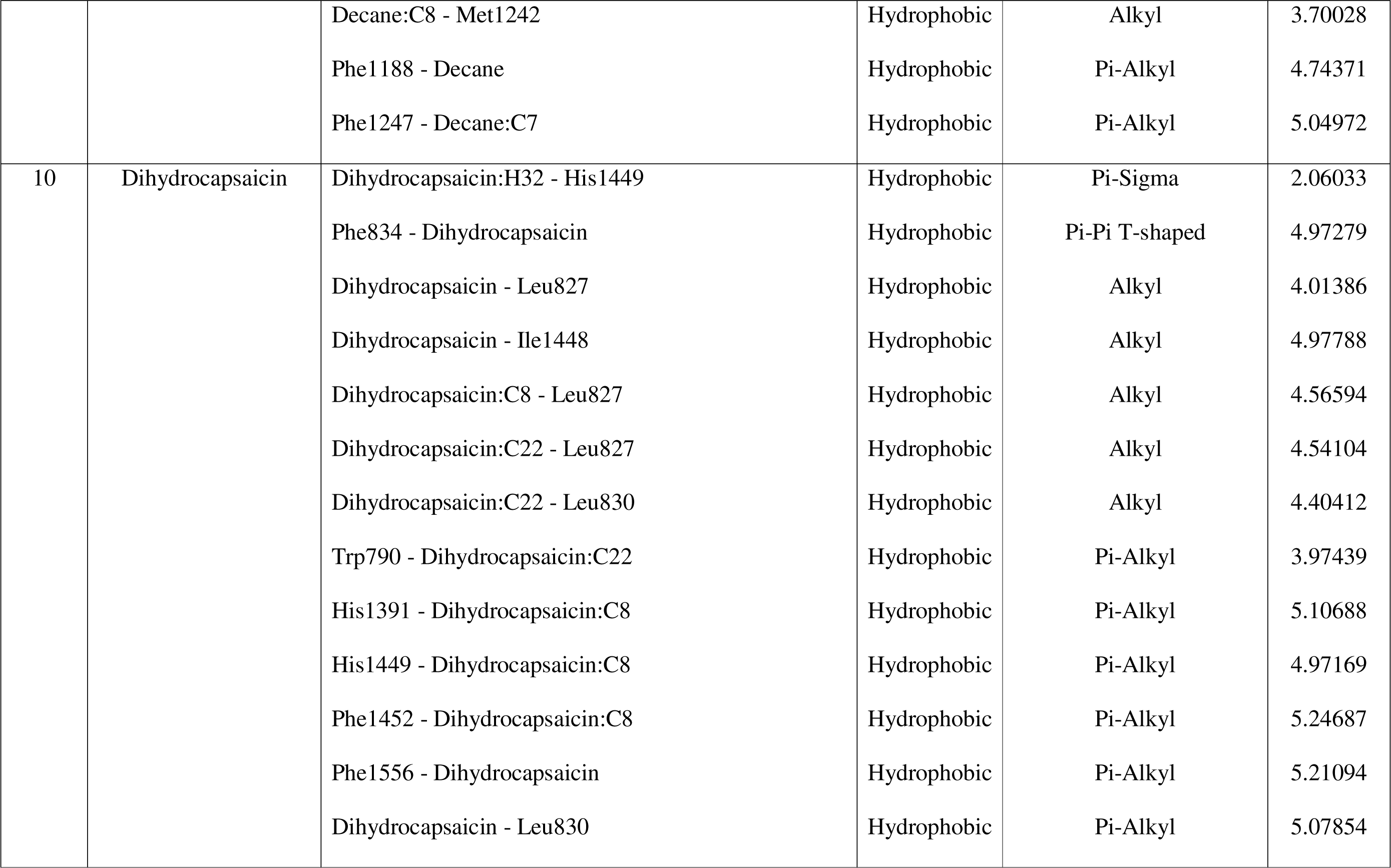
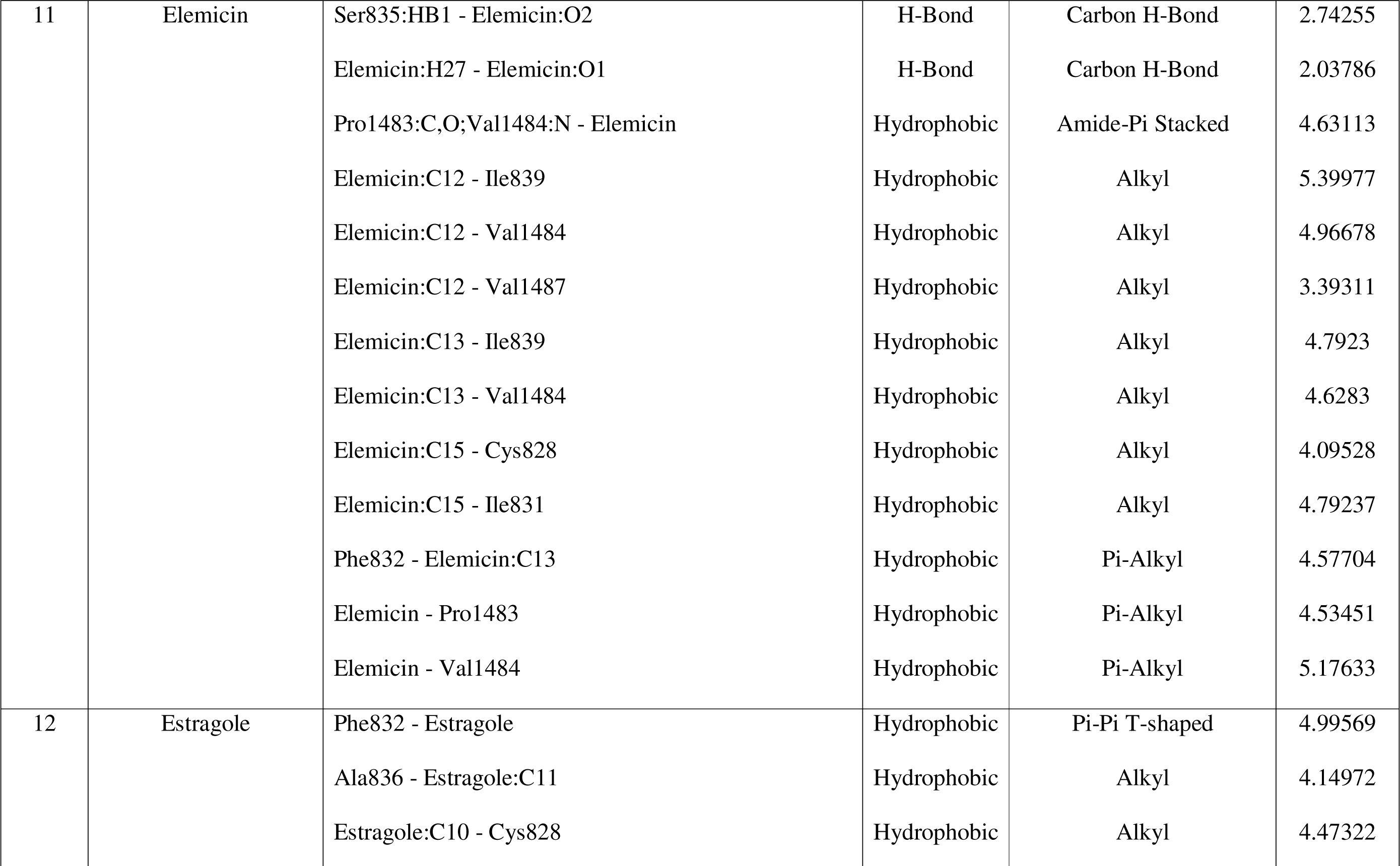
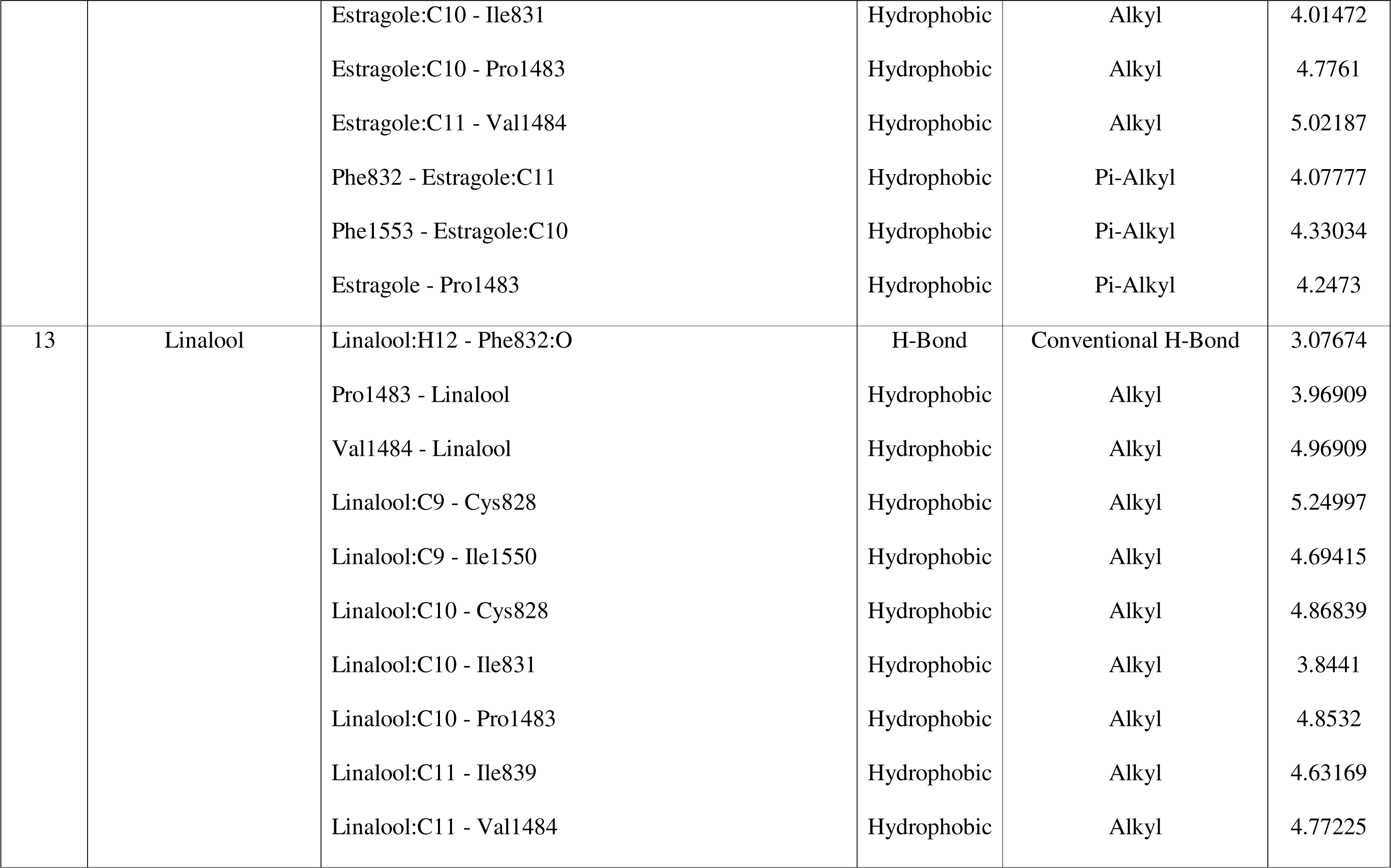
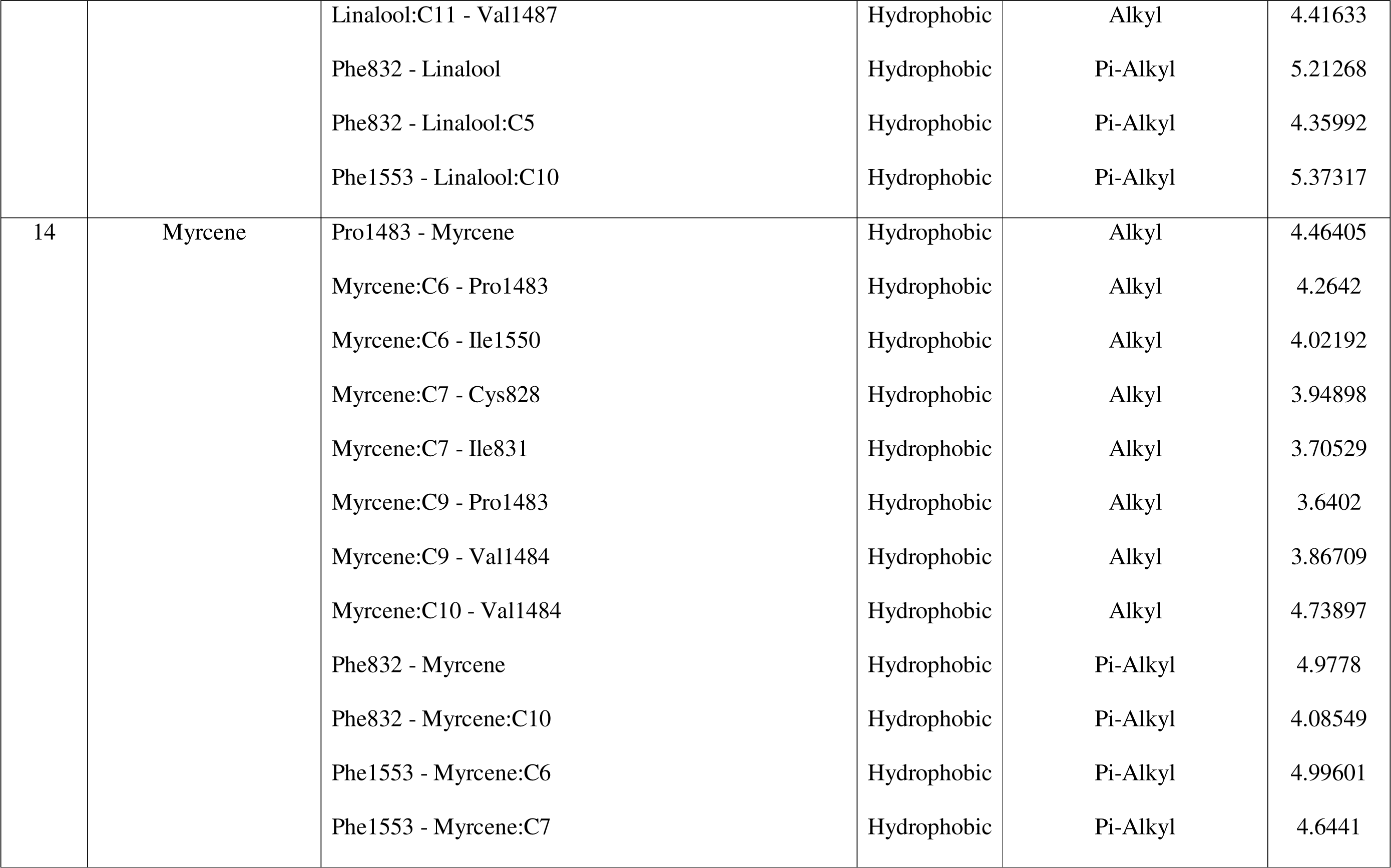
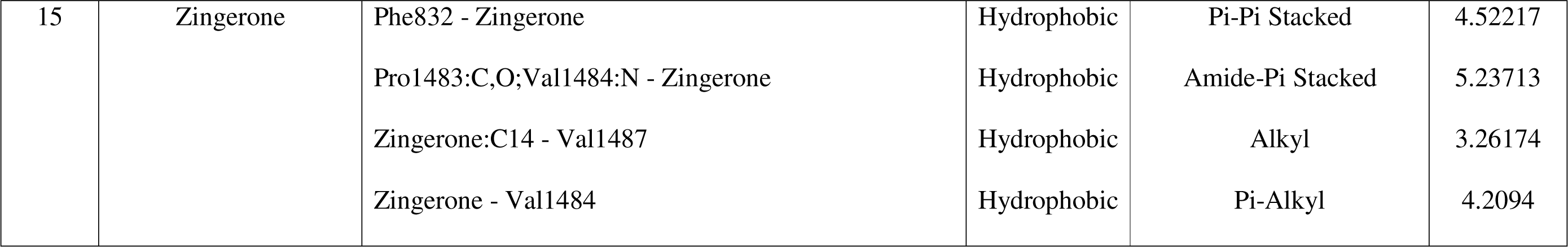
Molecular interactions of top 15 ligands.

## CONCLUSIONS AND FUTURE PERSPECTIVES

A total of 229 ligands (5 approved anti-fungal drugs, 10 x 5 = 50 drug derivatives, and 174 phytochemicals from IMPPAT database, with anti-fungal potential were considered for docking. The Chitin Synthase enzyme of *Rhizopus delemar* fungi was used as the key target to check the action of these ligands on this protein, computationally. As the 3D structure of the protein was not available in the PDB database, the protein was modelled using its amino acid sequence derived from NCBI. The model which was created using SWISS-DOCK was used as the final model for docking after Ramachandran plot validation and other parameters. Molecular docking was carried out using Discovery Studio Visualizer in order to investigate the binding affinity of these ligands, as mentioned above. All the ligands were tested for their compliance with the Lipinski’s rule, Veber’s rule, along with ADMET and TOPKAT parameters. Through docking it was found that Posaconazole 1 was the highest scoring ligand amongst the drug/drug derivatives with a LibDock score of 107.862. Among the phytochemicals, 1-nonacosanol emerged as the highest scoring ligand with a LibDock score of 118.483. The key residues in the protein interacting with the ligands were identified as Ile 831, Cys828, Phe832, Pro1483, Val1484. Out of the 25 ligands out of 229 which were able to bind to the protein, only 18 were able to satisfy the ADMET parameters and in addition to that, only 15 complied with the Lipinski’s rule and Veber’s rule. Dihydrocapsaicin was found to be the best ligand out of the final 15, considering all the parameters, such as Binding score, ADMET, TOPKAT, compliance to Lipinski’s and Veber’s rule. Cubebin was also shown to be a good lead molecule, but reported severe ocular irritancy alone as a disadvantageous effect. Thus, this study enables us to understand the interaction of various drugs/their derivatives and anti-fungal phytochemicals against the modelled protein hence, setting direction for designing better drugs against Mucormycosis by targeting its Chitin synthase enzyme.

Future prospects include wet-lab (in-vitro) studies using the phytochemicals and the drug derivatives; i.e., preliminary screening of the molecules identified using docking studies against the pathogen. The pathogen has to be isolated and identified, the phytochemicals need to be isolated from their plant sources and identified, modified drug derivatives need to be chemically synthesized, anti-fungal activity evaluation need to be performed using agar well diffusion method, MIC and IC50 determination for the lead phytochemicals, Chitin biosynthesis inhibition assay, and cytotoxicity testing.

